# Coenzyme A is a redox sensing cofactor for malic enzyme 2 regulating oxidative stress and mitochondrial metabolism

**DOI:** 10.64898/2026.04.27.721221

**Authors:** Wenzhe Chen, Ornella D. Nelson, Xiaorong Li, Manfeng Zhang, Xuan Lu, Tao Yu, Cong-Hui Yao, Shuai Zhang, Yugang Zhang, Adam Francisco, Byunghyun Ahn, Em Lundberg, Minsu Song, Janane F. Rahbani, Zhongzhou Chen, Hening Lin

## Abstract

Coenzyme A (CoA) is an essential cofactor required for numerous metabolic reactions, yet its ability to bind and regulate proteins remains poorly defined. Using a proteomic approach, we identified malic enzyme 2 (ME2) as a CoA-binding protein. ME2 uses NAD(P)^+^ to convert malate to pyruvate, generating NAD(P)H to support energy production and redox homeostasis. ME2 binds CoA at an allosteric site previously thought to bind NAD(P)^+^. Reduced CoA has minimal effect on ME2 activity, but the oxidized form, CoA disulfide, strongly activates ME2 by promoting ME2 tetramerization and a catalytically efficient closed conformation. Under oxidative stress, ME2 facilitates CoA disulfide formation, enhancing NADPH production and cellular defense against reactive oxygen species (ROS). Mice with ME2 mutation that cannot bind CoA show impaired muscle performance, elevated ROS, and mitochondrial dysfunction. These findings establish CoA as a redox-sensing cofactor, allowing cells to respond to ROS and promote mitochondrial metabolism, and expanding the function of this essential cofactor.

## Introduction

Coenzyme A (CoA), discovered by Fritz Lipmann in the 1940s ^1^, is a central metabolic cofactor that carries acyl groups via a high-energy thioester bond. It is required for mitochondrial energy production, fatty acid synthesis and degradation, cholesterol biosynthesis, and acyl-CoA–dependent protein modifications, including lysine acetylation, cysteine palmitoylation, and N-terminal glycine myristoylation.^2^ CoA has also been proposed to participate in redox signaling through protein CoAlation.^3, 4^ Given its pivotal role in cellular metabolism, disruption of CoA homeostasis has been implicated in numerous disease states, including cancer, metabolic disorders, and conditions associated with mitochondrial dysfunction and oxidative stress.^5^ Many important metabolites, including ATP and NADP^+^, are known to bind and regulate the functions of proteins ^6,7^. Despite the importance of CoA, only a limited number of proteins—such as pantothenate kinase (PanK) ^8, 9^ and calcium/calmodulin-dependent protein kinase II (CaMKII) ^10^—have been identified as being directly regulated by CoA, highlighting a significant gap in our understanding of its broader regulatory functions in biology.

To systematically identify CoA-binding proteins, we employed a biotinylated CoA probe (CoA–biotin) to enrich and identify proteins that directly interact with CoA. Using this approach, we identified malic enzyme 2 (ME2) as a previously unrecognized CoA-binding protein. ME2 is a mitochondrial enzyme that catalyzes the oxidative decarboxylation of malate to pyruvate, at the same time converting NAD^+^ or NADP^+^ to NADH or NADPH. This activity positions ME2 at a critical metabolic junction connecting different metabolic intermediates in the tricarboxylic acid cycle ^11–13^. Through this activity, ME2 contributes to cellular energy production via NADH generation and plays an important role in redox homeostasis by supporting NADPH availability ^14–16^.

Interestingly, we found that ME2 directly binds CoA and utilizes CoA to sense the redox state and tune its enzymatic activity. This reveals a previously unrecognized function of CoA in cellular defense against reactive oxygen species (ROS). Furthermore, mice harboring mutations at the ME2 CoA binding site exhibit significantly impaired muscle performance in treadmill running tests. This phenotype is accompanied by elevated mitochondrial ROS and diminished mitochondrial metabolism. Together, these findings suggest the ME2-CoA interaction as a cellular redox sensor and a critical regulatory mechanism to ensure physiological homeostasis and fitness.

## Results

### Identification of ME2 as a CoA-binding protein using Biotin-CoA

We synthesized a Biotin-CoA probe (Figure 1A) to pull down potential CoA-binding proteins. To account for background interactions caused by biotin or streptavidin beads, we used two control groups, one treated with Biotin-azide only, and one treated with excessive CoA as a competitor for Biotin-CoA binding (Figure 1A). Proteins enriched exclusively in the Biotin-CoA group were considered true targets (depicted in pink in Figure 1A) and ranked based on the Heavy-to-light (H/L) ratio. Nucleoside diphosphate kinase A (NME1) and B (NME2), which were previously reported to interact with long-chain fatty acyl coenzyme A (LCFA-CoA) ^17^, were among the most enriched proteins. Another protein, Malic enzyme 2 (ME2), showed an even stronger enrichment for CoA binding (Figure 1B). ME2 is a critical mitochondrial malic enzyme that catalyzes the oxidative decarboxylation of malate to pyruvate while reducing NAD(P)^+^ to NAD(P)H, and has never been reported as a CoA-binding protein.

**Figure 1.**
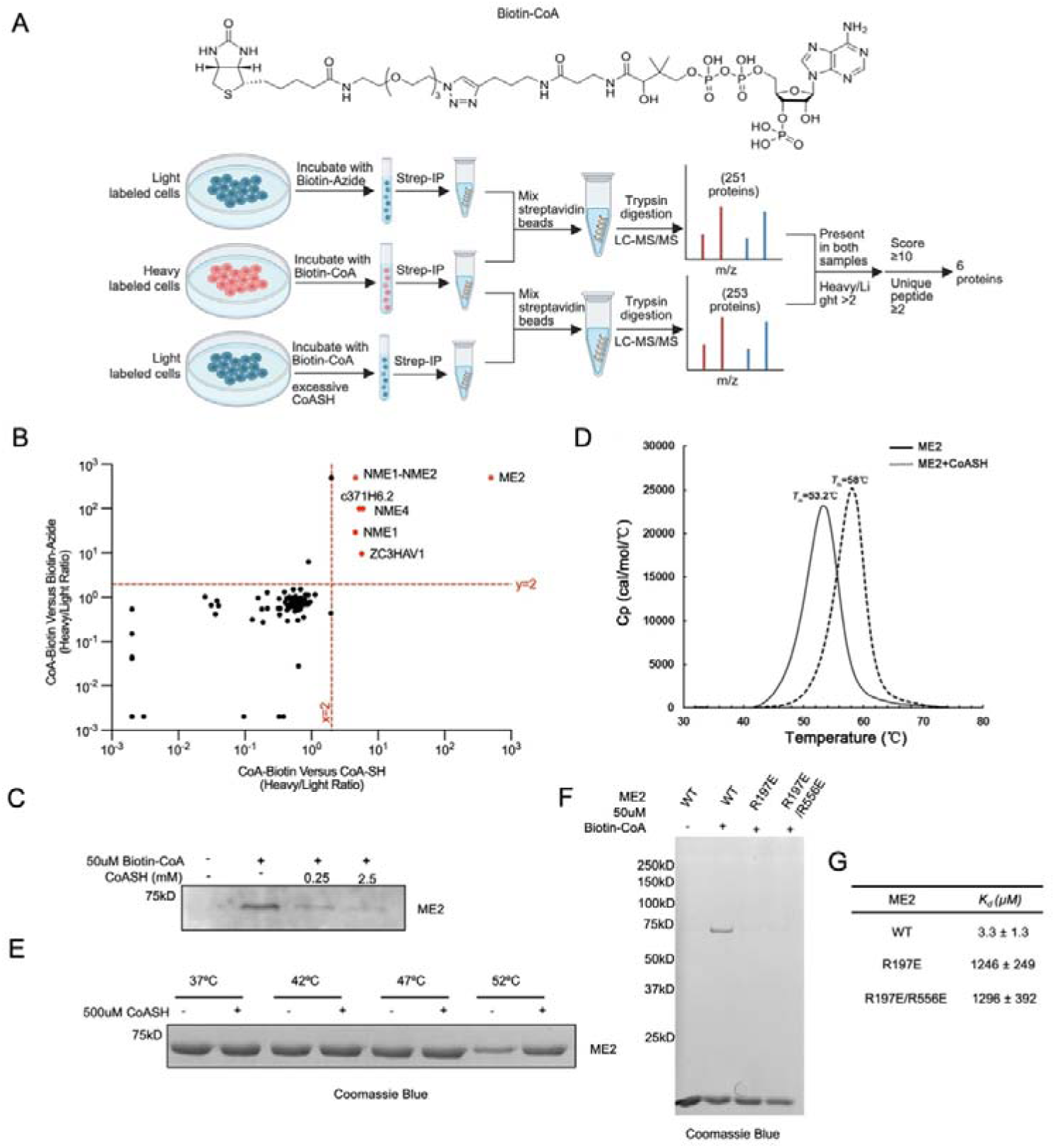
Identification of ME2 as a CoA-binding protein using Biotin-CoA. (A) SILAC proteomics was performed using Biotin-CoA with either Biotin-azide or CoASH competition. Scheme of data processing workflow to idenitfy Biotin-CoA specific binding proteins is shown. (B) Scatter plot showing all the hits from the proteomic analysis. The hits passing the threshold (Heavy/Light ratio >2) were labeled in red and the rest were labeled in black. (C) ME2 protein in cells was enriched using Biotin-CoA and CoASH was able to compete with binding. (D) Representative melting curves of ME2 and ME2-CoASH complex. The peak maxima of the derivative curves correspond to the protein melting temperatures (*T*_m_). (E) In-gel thermoshift assay was performed to verify the binding between purified ME2 protein and CoASH. (F) Purified ME2 WT proteins can be enriched using Biotin-CoA but not the exo site mutants. (G) Binding affinity CoASH to WT and exo site mutants.

To validate ME2 as a CoA-binding protein, we ran an in-vitro competition assay with Biotin-CoA and reduced free CoA (CoASH) using cell lysate. Our results show that endogenous ME2 can be enriched by Biotin-CoA, and excess CoASH competes with this interaction (Figure 1C). Using purified recombinant human ME2 protein, differential scanning calorimetry (DSC) experiments showed that the addition of CoASH increased the T_m_ value of ME2 by about 5 degrees, and the melting curve became more uniform (Figure 1D). We confirmed this effect using an gel-based thermal shift assay (Figure 1E). These results indicate that CoASH can bind and improve the thermal stability of ME2.

We next sought to identify the potential CoA binding site. Given that NAD^+^ and CoA have similar structures, we reasoned that CoA could bind to the same pockets that accommodate NAD^+^, which is known to bind ME2 at both the active site and an allosteric ‘exo’ site ^18,19^. Because CoA is unlikely to be the substrate of ME2, we hypothesized that it binds to the exo site. To test this hypothesis, we mutated two important arginine residues at the human ME2 exo site, R197E and R556E. Indeed, purified ME2 enzyme containing these mutations were not enriched by Biotin-CoA, suggesting that the exo site is the binding pocket for CoA (Figure 1F). We also measured the binding affinity of CoASH to WT and mutant ME2 proteins. WT ME2 binds to CoASH with a KD of 3.3 µM, while the R197E and R197E/R556E mutants have dramatically decreased binding affinities to CoASH (Figure 1G).

### CoA disulfide activates ME2 enzyme activity

We next investigated how CoA binding affects ME2 enzymatic activity. We monitored purified recombinant ME2 activity by measuring the absorbance of NAD(P)H at 340 nm in prescence of CoASH and several common CoA species, including acetyl-CoA, propionyl-CoA, butyryl-CoA, succinyl-CoA and malonyl-CoA. Unexpectedly, ME2’s enzymatic activity was minimally affected by these cofactors, and the largest effect was observed for CoASH, which increased ME2’s activity by just ∼1.2 fold.

To understand why CoASH binding would have such a modest effect on ME2’s enzymatic activity, we examined ME2’s previously solved X-ray crystal structure, which reveals that the active ME2 tetramer has two exo sites adjacent to each other at the tetramer interface ^20^. If each exo site binds to CoA, the adjacent CoA molecules could potentially dimerize to form CoA disulfide under oxidative stress, and bind ME2 more tightly to promote active tetramer formation. In this scenario, CoA disulfide may activate ME2 activity better than CoASH. Strikingly, the CoA disulfide (100 µM) shows a much stronger (∼2-fold) effect on ME2 enzyme activity (Figure 2A). This effect is concentration-dependent: even 0.1 uM of CoA disulfide activated ME2 dramatically, and 10 μM CoA disulfide increased its activity by almost 10-fold (Figure 2B). However, 100 µM CoA disulfide showed only ∼2-fold ME2 activation. This is likely due to the hook effect - with excessive CoA disulfide, each ME2 monomer will bind one CoA disulfide, decreasing ME2 tetramer formation (tetramer formation requires one CoA disulfide bound by two ME2 monomers). Next, we compared CoA disulfide with fumarate, which is known to activate ME2 ^13^. We observed that CoA disulfide (10 μM) activates ME2 several fold better than fumarate (at a previously established concentration of 4 mM), making CoA disulfide the strongest ME2 activator discovered so far (Figure 2C).

**Figure 2.**
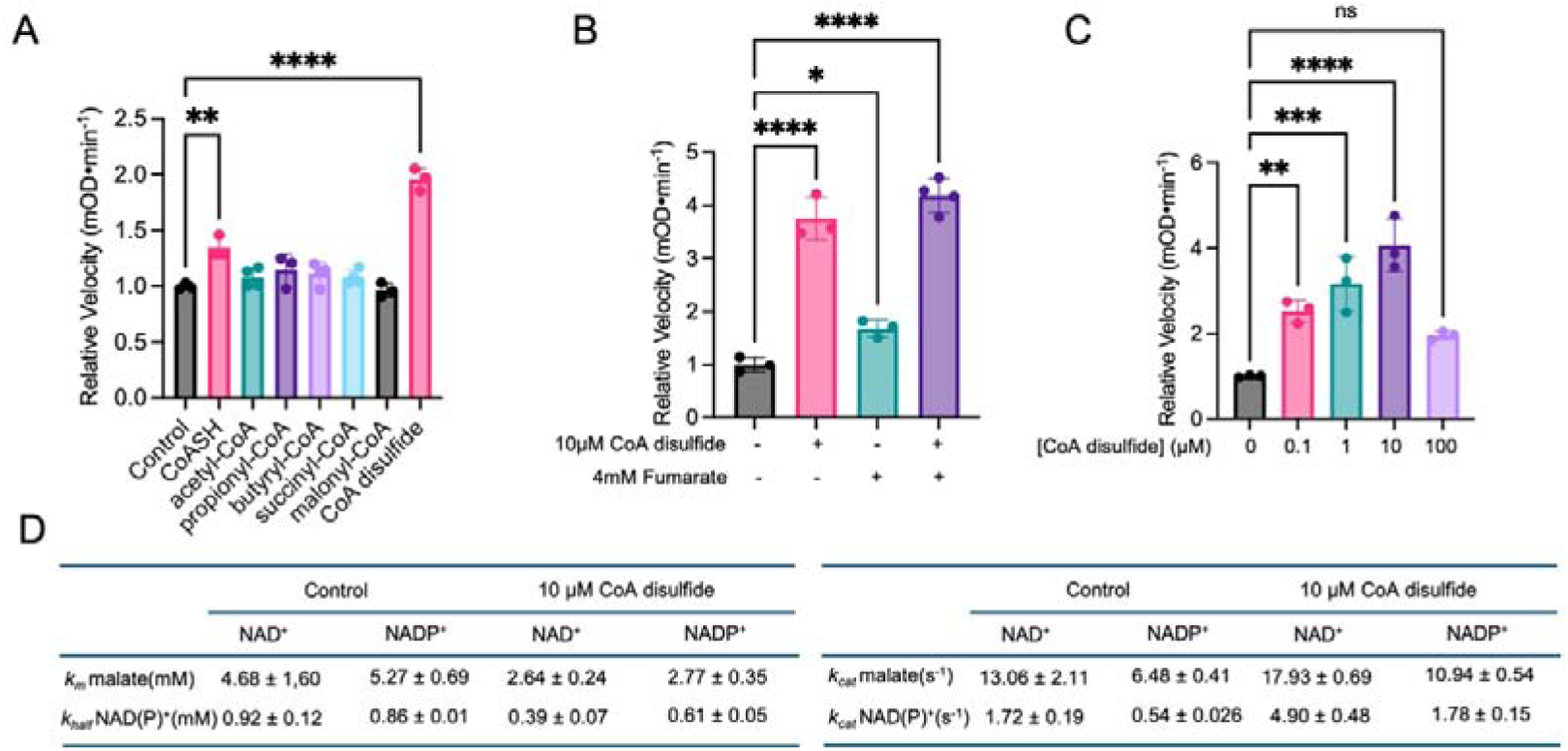
CoA disulfide activates ME2 enzymatic activity. (A) Various CoA species were screened in vitro, and CoA disulfide was found to be the strongest ME2 enzyme activator. (B) CoA disulfide was compared with previously known ME2 activator, fumarate, and CoA disulfide showed better ME2 activation in vitro. (C) Several CoA disulfide concentrations were tested on ME2 enzymatic activity. (D) Kinetic parameters for NAD^+^ and NADP^+^ under conditions without and with 10 µM CoA disulfide. The results in (A), (B) and (C) are shown as mean ± SD. ns, not significant. *P < 0.05; **P < 0.02; ***P < 0.002; ****P < 0.001.

We then performed more detailed enzyme kinetic studies of ME2 with or without 10 µM CoA disulfide (the *K_m_* and *k_cat_* of malate and NAD(P)^+^ are summarized in Figure 2D and plotted in Figure S1A-C). CoA disulfide decreased the *K_m_* and increased the *k_cat_*, regardless of whether NAD^+^ or NADP^+^ was used as the cofactor.

### CoA disulfide promotes ME2 tetramer formation by binding to the exo site

To determine whether CoA disulfide binding could promote and stabilize the ME2 active tetramer, we used size exclusion chromatography (SEC) to examine the size distribution of ME2 in the presence of different ligands. Under our experimental conditions, the inactive apo-ME2 primarily exists as a dimer in solution, with the major peak exhibiting a volume value of 15.5 mL. Fumarate and NAD^+^ have no measurable effect on the size distribution of ME2. In contrast, the binding of CoASH or CoA disulfide allows ME2 to exist mainly as a tetramer, with a peak volume value of 13.2 mL (Figure 3A). We also used SAXS, a powerful tool for the quantitative analysis of the aggregation state in a solution, to show that the binding of CoA or disulfide-CoA transformed ME2 from mainly a dimer into mainly a tetramer (Figure 3B, C, D; Supplementary Table 1).

**Figure 3.**
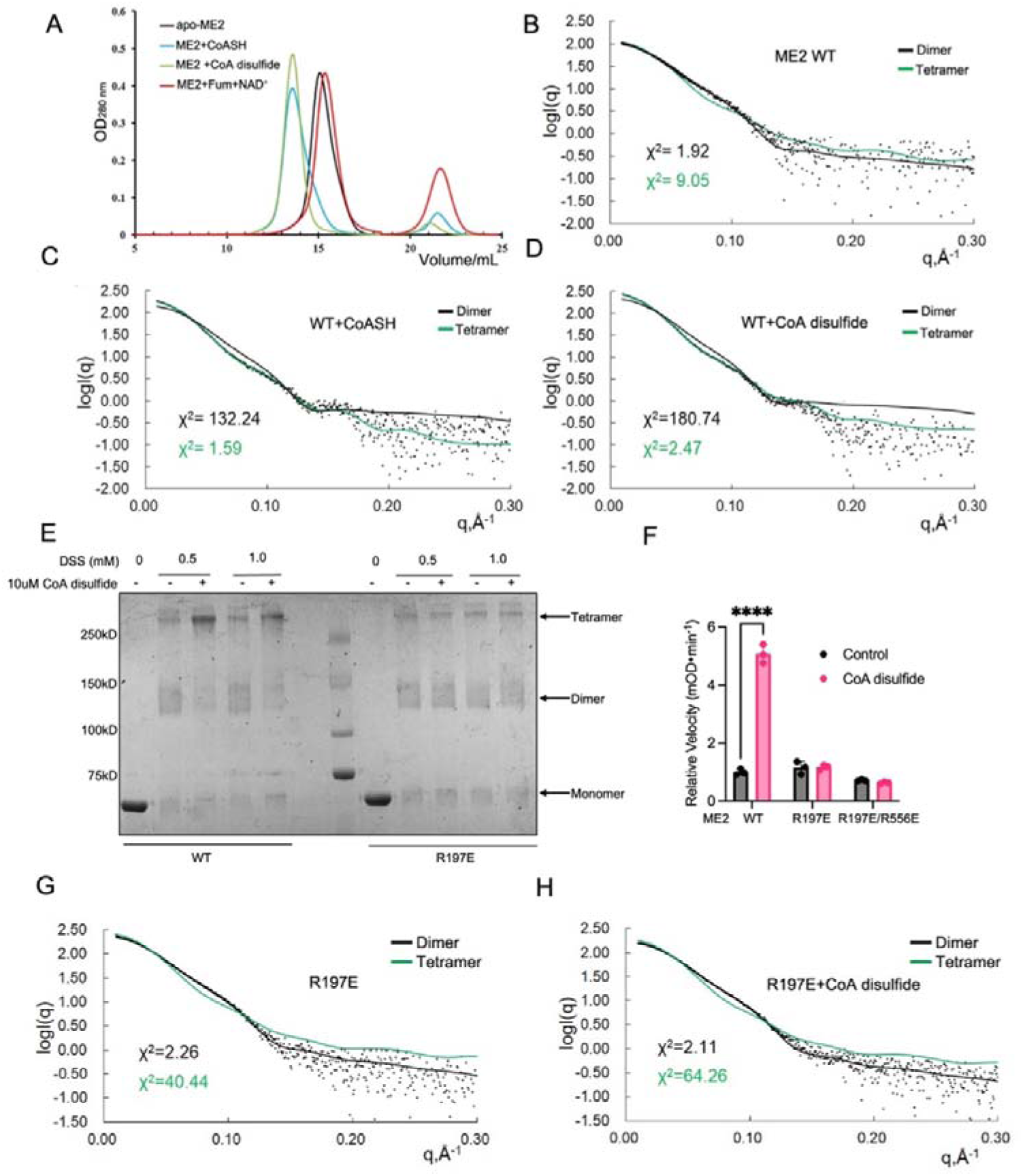
CoA disulfide promotes ME2 tetramer formation. (A) Size exclusion chromatography profiles of 2.5 mg/mL ME2 WT on Superdex 200 10/300 GL column, alone and in the presence of 1 mM CoASH, Fumarate/NAD^+^, or CoA disulfide. (B-D) Overlay of experimental scattering profiles with calculated scattering profiles for 1 mg/mL ME2 WT (B), ME2 WT with CoASH (C), and ME2 WT with CoA disulfide (D). (E) Detecting the oligomeric state of ME2 using SDS-PAGE after crosslinking with DSS. CoA disulfide promotes tetramer formation in ME2 WT, but not in the exo site R197E mutant. (F) Activity of ME2 WT and mutant with saturated malate *in vitro* (means ± SD, n = 3). (G, H) Overlay of experimental scattering profiles with calculated scattering profiles for R197E alone (G) and with CoA disulfide (H). Experimental data are represented in black dots. The theoretical scattering curves of tetramers (green lines) and dimers (black lines) are shown. Dimer is constructed based on two protomers located on both sides of the dimer interface. The results in (F) are shown as mean ± SD. ns, not significant. *P < 0.05; **P < 0.02; ***P < 0.002; ****P < 0.001.

To visualize the ME2 tetramer promoted by CoA disulfide on an SDS-PAGE gel, we used the chemical crosslinker, disuccinimidyl suberate (DSS), to crosslink purified ME2 WT or the exo site R197E mutant, with or without CoA disulfide. CoA disulfide promoted the tetramer formation of ME2 WT but not the R197E mutant (Figure 3E). Additionally, we found that CoA disulfide can only activate ME2 WT, but not the R197E mutant, even though they have similar basal enzyme activity without CoA disulfide (Figure 3F). Using SAXS, we further validated that CoA disulfide only promotes tetramer formation for ME2 WT but not R197E mutant enzyme (Figure 3G, H).

### X-ray crystal structures clarify the interaction of ME2 in complex with CoASH versus disulfide-CoA

To determine the structural basis of ME2 regulation by CoA and disulfide CoA, we solved four crystal structures of human ME2 in two binary complexes with CoASH or CoA disulfide, as well as two quinary complexes with CoASH or CoA disulfide, NAD^+^, pyruvate (Pyr), and Mn^2+^ (statistics for diffraction data collection and structure refinement are summarized in Supplementary Table 2). All the protomers in each structure have similar conformations, with a root-mean-square deviation (r.m.s.d.) distance of ∼1 Å between all equivalent Cα atoms. Additionally, we obtained the structure of the apo form of ME2 without any ligands bound. The overall fold of the apo-ME2 is similar to other structures, but with higher flexibility. The protomer conformation and tetramer organization of apo-ME2 represent the original open form I state. Indeed, as previously hypothesized, CoASH and CoA disulfide bound to the exo site of ME2 located at the tetramer interface (Figure 4A) ^19,20^. The ME2 tetramer binds either four CoASH or two CoA disulfide molecules.

**Figure 4.**
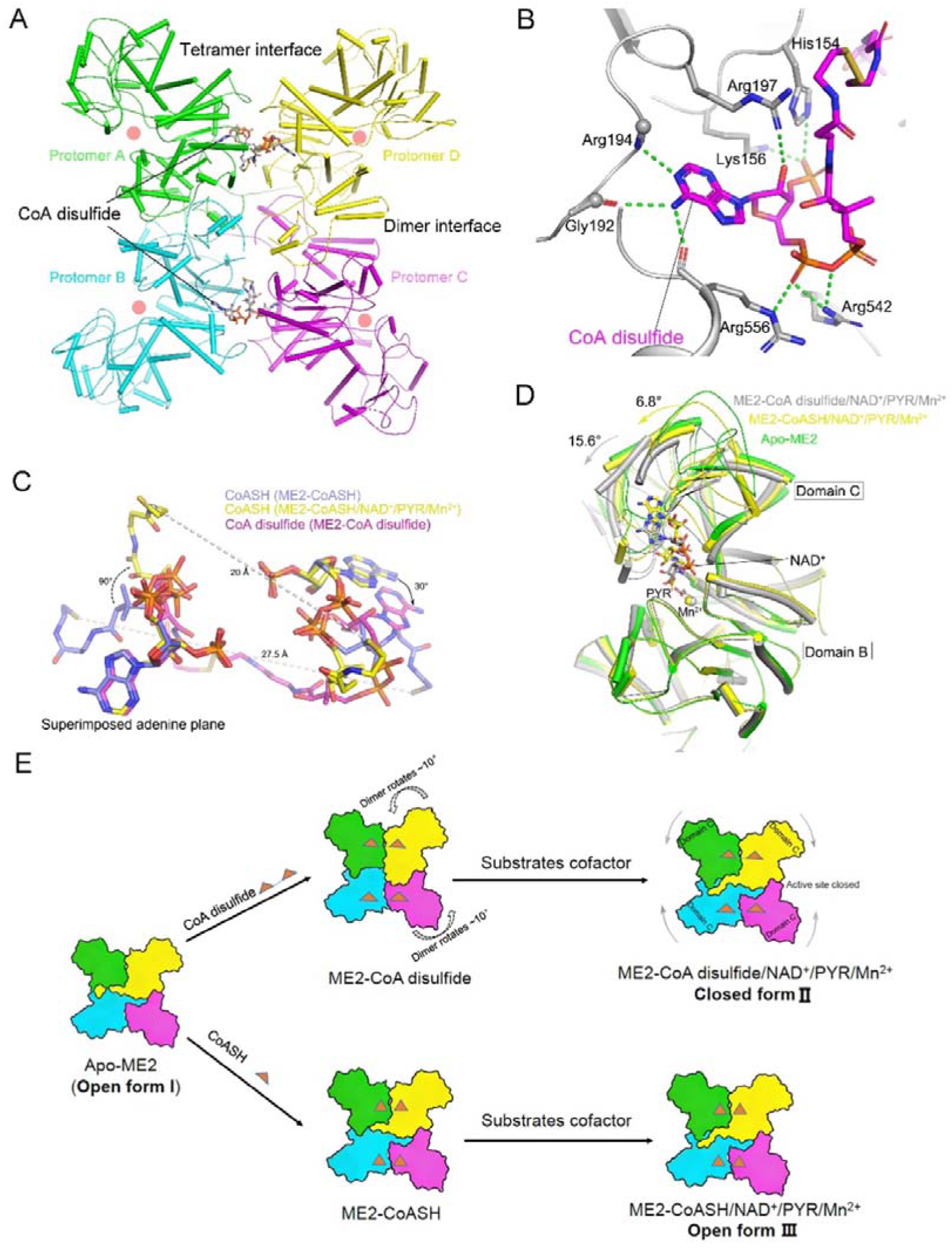
Structural analysis of ME2 in complex with CoA disulfide and comparisons with other ME2 complexes. (A) The tetrameric structure of ME2 in complex with CoA disulfide. The protomers are colored in green, cyan, yellow and purple. The two CoA disulfide molecules are shown as stick models colored according to atom type. The dimer and tetramer interfaces are labeled, and the active site is highlighted by the light pink circle. (B) Diagram of the binding site of CoA disulfide in the ME2-CoA disulfide complex structure. CoA disulfide (magenta) and interacting protein residues (gray) are shown as sticks. Hydrogen bonds between the protein and the cofactor are indicated as green dotted lines. (C) Schematic drawing showing the comparison of molecular conformation of CoASH (slate and yellow represent ME2-CoASH binary and ME2-CoASH/NAD^+^/PYR/Mn^2+^ quinary complex structures, respectively) and CoA disulfide (purple). (D) Protomer superposition of apo-ME2, ME2-CoASH/NAD^+^/PYR/Mn^2+^, and ME2-CoA disulfide/NAD^+^/PYR/Mn^2+^ structures. (E) Scheme summarizing the structural changes of ME2 tetramers due to binding of substrates and cofactors. Only CoA disulfide induces a profound conformational reorganization.

In all the ME2 complex structures, the electron density of the 3’-phospho-ADP portion of CoASH or CoA disulfide is clearly visible at the “exo” site (Figure S2A, S2B). As expected, the adenine base and ribose part of CoASH or CoA disulfide interacts with ME2 similarly to NAD^+^ and ATP (Figure S2C). The N6 amino group of the adenine base forms two hydrogen bonds with the main chain carbonyls of Arg556 and Gly192, and the N1 ring nitrogen makes a hydrogen bond to the main chain amide of Arg194, supporting the recognition of the adenine base. The Arg197 side chain is involved in forming a large, positively charged surface and binds CoASH through hydrogen bonds, electrostatic interactions, and amino-aromatic interactions (Figure 4B, S2D-2F). The 3’-phosphate group of ribose forms interactions with the side chains of Lys156 and His154. The α-phosphate interacts with the side chains of Arg542 and Arg556 (Figure 4B, S2D, S2E). The rest of the CoASH or CoA disulfide molecule makes a few contacts with ME2. The steric hindrance caused by the 3’-phosphate and β-mercaptoethylamine groups may explain the specificity of CoASH or CoA disulfide for the exo site over the active site. We mutated relevant residues and used SAXS to detect the oligomeric state of ME2 in solution upon CoA disulfide binding. As mentioned earlier, the most significant effect was observed with the R197E mutation (Figures 3G, 3H, S3A-F).

### CoA disulfide induces a profound reorganization of the ME2 tetramer to form a catalytically efficient closed conformation

At the tetramer interface, the relative positions of two CoASH molecules and one CoA disulfide molecule exhibit significant variability. In the ME2-CoASH binary complex, CoASH adopts a "U" shape, with the pantotheine and β-mercaptoethylamine groups arranged anti-parallel to those of adjacent CoASH molecules at the interface (Figure 4C). Upon NAD^+^/PYR/Mn^2+^ binding, both groups rotate ∼90° in the same direction (Figure 4C). These structural changes bring the two sulfhydryl groups closer together and orient them toward the tetramer surface. In the ME2-CoA disulfide binary complex, the disulfide bond increases interconnectivity between the bound protomers. When one adenine plane is aligned, the other rotates ∼30° compared with the CoASH-bound structure (Figure 4C). Since each CoA disulfide bridges two protomers across the tetramer interface, this binding mode induces relative rotation between them, ultimately driving reorganization of the tetramer. Thus, the conformational transition from CoASH to CoA disulfide directly alters the tetramer organization of ME2.

Each protomer of the ME2 tetramer can be divided into four domains (A–D; Figure S3G). Careful analysis reveals marked differences in the conformations of the protomers and tetramers across these structures. When domains A and B are aligned, domain C of the ME2-CoA disulfide/NAD^+^/PYR/Mn²□ complex requires a 15.6° rotation to superimpose with that of apo-ME2 (Figure 4D), whereas a 6.8° rotation is needed for the ME2-CoASH/NAD+/PYR/Mn²□ complex to superimpose with that of apo-ME2. The quinary ME2-CoA disulfide/NAD+/PYR/Mn²□ complex adopts a closed conformation, consistent with previously reported ME2-substrate complex structures. Superposition of 548 equivalent Cα atoms from protomers of the ME2-CoA disulfide/NAD^+^/PYR/Mn²□ complex and the closed-form structure (PDB: 1PJ2) yields a r.m.s.d. of 0.4 Å. In contrast, the quinary ME2-CoASH/NAD^+^/PYR/Mn²□ complex remains in an open conformation. Both the bindings of CoASH and CoA disulfide to ME2 promote the formation of more ME2 tetramers (Figures 3B-D). However, the CoA disulfide-bound ME2 tetramer undergoes substantial conformational reorganization, assuming a tetrameric state closer to previously reported closed conformation. Subsequent substrate binding further drives this CoA disulfide-bound tetramer into a fully closed state (II), which may underlie the allosteric activation of ME2 by CoA disulfide. Conversely, the CoASH-bound ME2 tetramer conformation (III) more closely resembles the open conformation of apo-ME2 (I). Even upon substrate binding, it retains a conformation near the open state (Figure 4E). The binding of pyruvate in this state (III) differs distinctly from that in the closed state (II). In state III, water mediates the interaction between Mn^2+^ cation and pyruvate while in the closed state II, pyruvate directly interacts with Mn^2+^ (Figures S3H and S3I). Collectively, CoA disulfide binding induces extensive conformational reorganization of the ME2 tetramer into a catalytically efficient near-closed state, consistent with its stronger potentiation of enzymatic activity.

### ME2 facilitates CoA disulfide formation under oxidative stress and CoA disulfide activates ME2 to reduce oxidative stress in cells

CoA and CoA disulfide were previously reported as important redox regulators in certain prokaryotic cells, analogous to the GSH/GSSG system in mammalian cells.^21–23^ Furthermore, recent studies suggest that CoA disulfide is a potential product of oxidative stress in mammalian cells, and may function as an intermediate metabolite promoting protein CoAlation.^3,21,24^ Although no direct evidence of CoA disulfide has been reported in mammalian cells, it is reasonable for two CoASH molecules to form a disulfide bond under oxidative stress, especially when they are in proximity, as when bound to ME2. Thus, we hypothesized that ME2 could facilitate the formation of CoA disulfide. After mixing purified ME2 with CoASH and using HPLC to analyze the level of CoASH and CoA disulfide, we indeed observed an increase in the ratio of CoA disulfide to CoA (Figure 5A). To further validate that ME2 promotes cellular CoA disulfide formation, we knocked down *ME2* in A549 cells and measured the cellular CoA disulfide/CoASH ratio using LC-MS. As expected, ME2 knockdown decreased the ratio of CoA disulfide to CoASH (Figure 5D).

**Figure 5.**
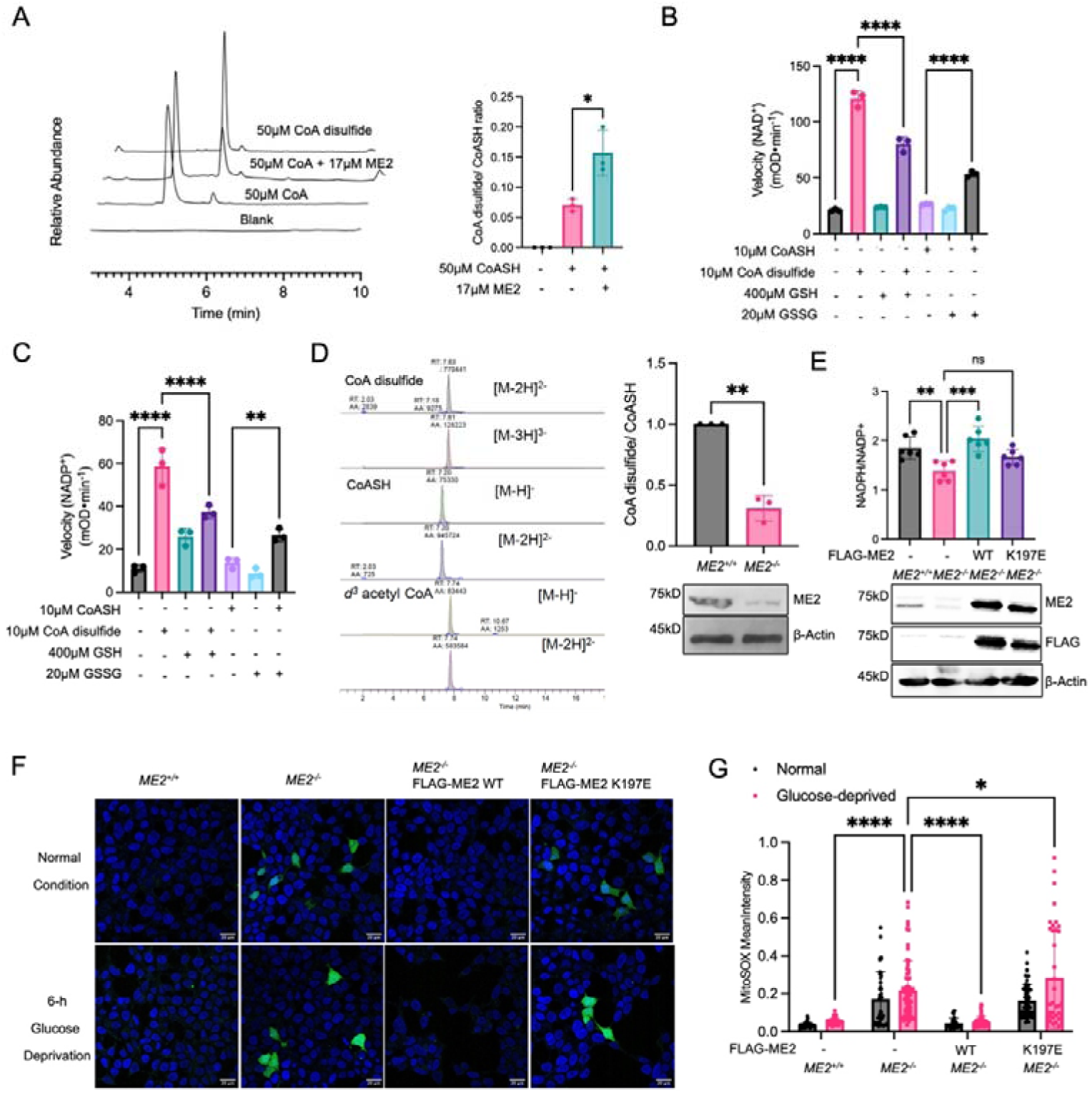
ME2 promotes CoA disulfide formation under oxidative stress and CoA disulfide activates ME2 to reduce oxidative stress in cells. (A) ME2 was mixed with CoASH and incubated at 37 °C for 30 mins. CoA disulfide/CoASH ratio was measured using HPLC and representative HPLC traces are shown. (B, C) ME2 activation by CoASH or CoA disulfide was measured under different redox conditions. The initial rate was measured with the addition of GSH and GSSG using either NAD^+^ (B) or NADP^+^ (C) as a substrate. The presence of GSH decreased ME2 activation and GSSG increased ME2 activation. (D) Cellular CoA disulfide/CoASH ratio was measured in A549 *ME2* knockdown cells using LC-MS. Representative LC-MS traces are shown on the left. (E) NADPH/NADP^+^ ratio was measured in HEK293T *ME2* KO cells re-expressing ME2 WT or R197E mutant. (F, G) ME2 WT or R197E mutant was reintroduced into the HEK293T *ME2* KO cells and stained with Mito-SOX probe. Represented images were shown (F) and quantified in (G). The results in (A-E) and (G) are shown as mean ± SD. ns, not significant. *P < 0.05; **P < 0.02; ***P < 0.002; ****P < 0.001.

Our results suggest a model in which ME2 uses the bound CoA molecule to sense oxidative stress and regulate its activity. Oxidative stress drives the formation of CoA disulfide on the ME2 exo site, which in turn activates ME2 to produce NADPH that is needed to remove oxidative stress.

To further evaluate this model, we performed ME2 activity assays in the presence of GSH and GSSG, which are the most abundant cellular redox species ^25^. We found that in the absence of CoASH or CoA disulfide, neither GSH nor GSSG had much effect on ME2 enzyme activity (Figure 5B, C). However, GSH decreases ME2 activation by CoA disulfide and GSSG increases ME2 activation by CoASH (Figure 5B, C), supporting that through CoASH and CoA disulfide, ME2 senses the redox state to tune its enzymatic activity.

To further test the model, we asked whether the exo site mutant R197E would have a measurable effect on the cellular NADPH/NADP^+^ ratio. We created a *ME2* HEK293T knock-out cell line. As expected, these cells exhibited a decreased ratio of NADPH to NADP^+^. We then re-expressed either WT ME2 or the R197E mutant in *ME2* knockout cells. WT ME2 brought the ratio of cellular NADPH/NADP^+^ back to normal, while the R197E mutant did not, suggesting that ME2 and its regulation by CoA play an important role in NADPH production (Figure 5E).

To further investigate the effect of ME2 and its regulation by CoA on cellular redox homeostasis, we measured levels of mitochondrial superoxide (MitoSOX), one of the major ROS in mitochondria, using a MitoSOX probe.^26^ We cultured cells in normal media and glucose starvation media, which increases oxidative stress. Consistent with the NADPH/NADP^+^ ratio result obtained above, *ME2* knockout cells had higher MitoSOX levels than *ME2* WT cells. In the *ME2* knockout cells, MitoSOX levels were rescued by ME2 WT re-expression, but not R197E mutant re-expression. While the MitoSOX was increased during glucose starvation, the overall trends were similar to that observed without glucose starvation (Figure 5F, G). Overall, these results support that ME2 uses CoA as a redox sensing cofactor to maintain cellular ROS homeostasis.

### *Me2 K197E* mutant mice exhibit impaired muscle function associated with mitochondrial dysfunction

As ME2 has been reported to play important roles in cancer and neurological diseases,^27–31^ we sought to investigate the physiological consequences of ME2 regulation by CoA and redox state. We initially tested the effect of the ME2 R197E mutation in several human cancer cell lines but did not observe overt proliferative or survival phenotypes (Figure S4). Although R197E disrupts the redox regulation of ME2, it does not affect ME2’s basal activity, potentially explaining the lack of a strong cancer phenotype under basal conditions.

To further explore its potential role in physiology, we generated Me2 transgenic mice in which the CoA binding residue, K197 (equivalent to R197 in human ME2), was mutated to glutamate using CRISPR-based editing. We compared the metabolism of *Me2 WT* and *K197E* mutant mice using a metabolic cage and found no significant differences (Figure S5). This prompted us to examine conditions where mice might be under high energy demand and oxidative stress. Skeletal muscle represents a highly ROS-sensitive tissue due to its substantial mitochondrial activity and ROS production during exercise.^32–35^ However, the role of Me2 in muscle physiology has not been previously reported. To investigate the physiological relevance of the CoA-mediated redox regulation of Me2 in vivo, we assessed muscle performance using a treadmill exhaustion assay, measuring running time and distance until exhaustion. Strikingly, male *Me2 K197E* mice exhibited a greater than 50% reduction in exhaustion run distance compared with WT C57BL/6 controls (Figure 6A, B). Although the phenotype was less pronounced in female mice, it remained statistically significant, underscoring an important role for the CoA-mediated redox regulation of Me2 activity.

**Figure 6.**
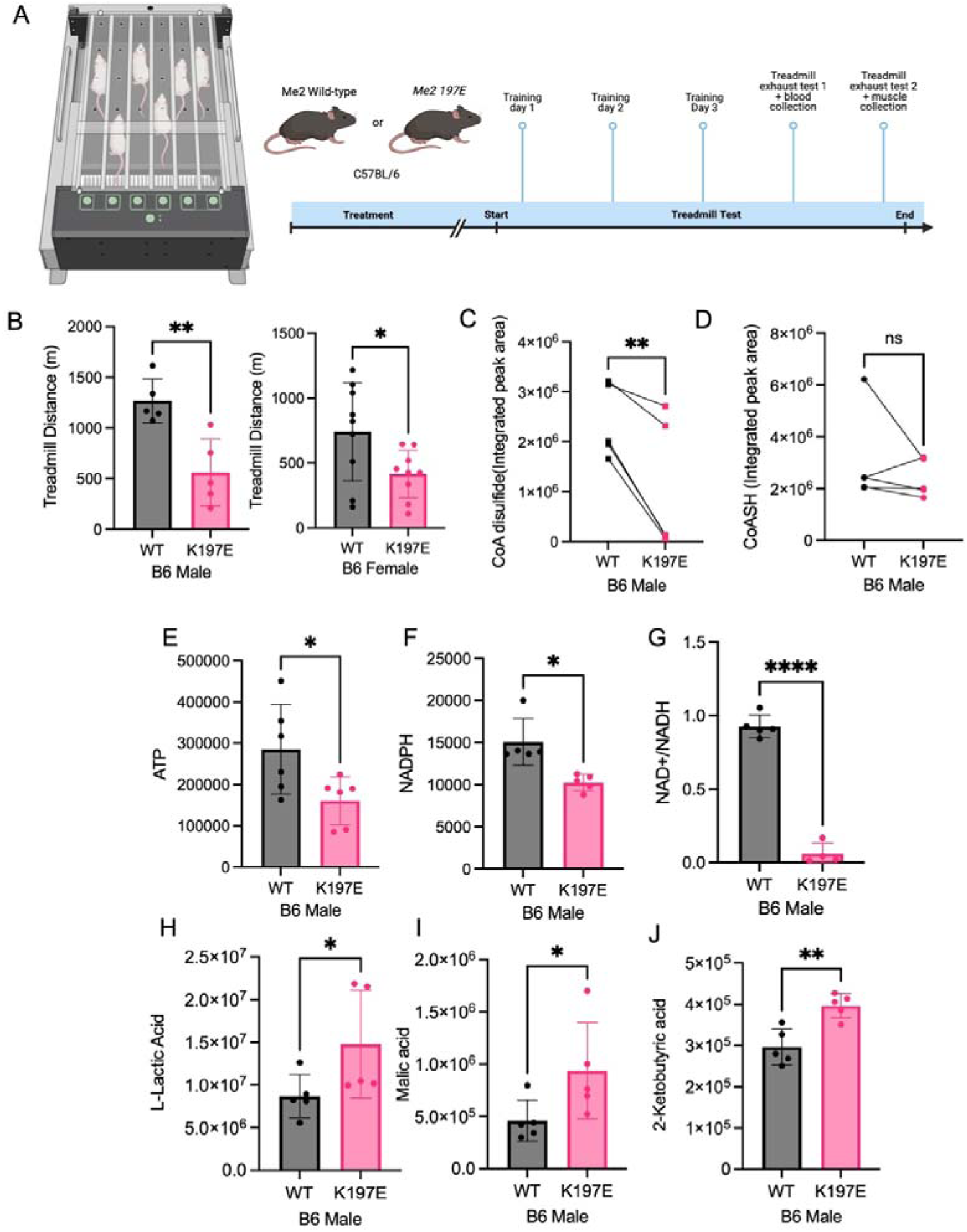
Me2 mutant mice have impaired muscle function and exercise capability due to mitochondrial dysfunction. (A) *Me2* WT and K197E C57BL/6J mice were subjected to the treadmill exhaustion test prior to the collection of muscle and blood samples. (B) The treadmill running distance until exhaustion was measured for both male and female mice. (C-G) Relative metabolite levels were measured in the muscle samples. C: CoA disulfide, D: CoASH, E: ATP, F: NADPH, G: NAD^+^/NADH ratio. (H-J) Relative blood metabolite levels in mice after treadmill exhaustion test. H: lactate, I: malate, and J: 2-ketobutyric acid. The results in (B-J) are shown as mean ± SD. ns, not significant. *P < 0.05; **P < 0.02; ***P < 0.002; ****P < 0.001.

**Figure 7.**
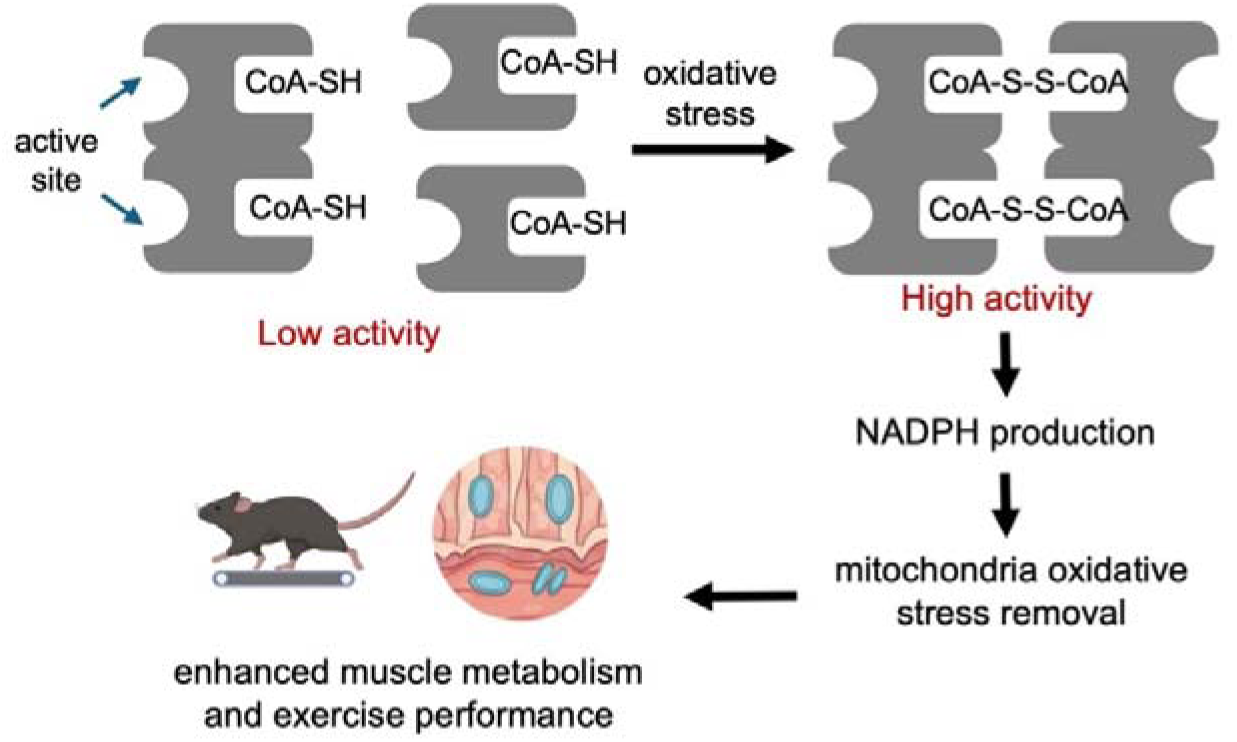
Proposed model of CoA-mediated redox regulation of ME2 that tunes ME2 activity to maintain redox homeostasis and mitochondrial metabolism. Under oxidative stress, ME2-bound CoASH forms CoA-disulfide, which strongly activates ME2 enzymatic activity and promotes NADPH production. The NADPH produced helps to remove oxidative stress and promote mitochondrial metabolism, leading to enhanced muscle metabolism and exercise performance.

To investigate the metabolic basis of this defect, mice were exercised to exhaustion and then sacrificed. We then collected muscle tissues and quantified different metabolites. Consistent with our earlier cellular data showing that ME2 knockout cells have decreased CoA disulfide, *Me2 K197E* muscle displayed a marked reduction in CoA disulfide levels, while CoASH levels remained largely unchanged (Figure 6C, D). These results from the *Me2 K197E* mice provide further support that Me2 uses CoA as a redox sensor to tune its activity to maintain redox homeostasis and mitochondrial metabolism. Consistent with impaired muscle performance, ATP levels were significantly reduced in *Me2 K197E* muscle compared with WT controls (Figure 6E). In addition, NADPH levels were decreased, consistent with increased oxidative stress and impaired mitochondrial redox homeostasis (Figure 6F). Notably, the NAD□/NADH ratio was dramatically reduced in *Me2 K197E* muscle (Figure 6G), suggesting a severe disruption in mitochondrial oxidative metabolism.

To further assess metabolic alterations at the systemic level, we analyzed serum metabolites following exercise. *Me2 K197E* mice exhibited elevated circulating lactate levels relative to WT mice (Figure 6H), indicative of increased reliance on glycolytic metabolism in the context of mitochondrial energy deficiency. Consistent with impaired TCA cycle flux, serum levels of TCA cycle intermediates, including malate and α-ketoglutarate, were significantly increased in *Me2 K197E* mice (Figure 6I, J). These data together suggest that loss of the CoA-mediated redox regulation of Me2 increases mitochondrial ROS during exercise, which compromises oxidative phosphorylation. As a consequence, NADH oxidation becomes inefficient, resulting in NADH accumulation and depletion of the NAD□ pool, thereby limiting ATP production and impairing muscle function.

## Discussion

Coenzyme A (CoA) is a central metabolic cofactor in mammalian cells. However, whether it can noncovalently bind to proteins and regulate protein function has not been well understood. Here, using SILAC-based proteomics combined with biochemical and structural validation, we identify mitochondrial malic enzyme 2 (ME2) as a novel CoA-binding protein, and CoA serves as a redox-sensing modulator of ME2 enzymatic activity. Under oxidative stress, the ME2 bound CoA forms CoA disulfide, which functions as a potent allosteric activator of ME2. We demonstrate that CoA species bind to a specific allosteric “exo” site that is distinct from the catalytic site and is essential for stabilizing the active tetrameric conformation of ME2. The dimeric CoA disulfide acts as a markedly strong activator, exceeding the effect of previously described activators such as fumarate by more than five-fold. Under oxidative stress conditions, ME2 facilitates the formation of CoA disulfide, which in turn enhances ME2 activity to promote NADPH production and maintain mitochondrial redox homeostasis.

The physiological relevance of this ME2–CoA axis is underscored by our *Me2 K197E* transgenic mouse model, in which disruption of CoA binding leads to a profound reduction in exercise capacity and severe mitochondrial dysfunction in skeletal muscle. *Me2 K197E* mice exhibit diminished ATP levels, elevated oxidative stress, and a dramatic imbalance in mitochondrial redox state, establishing the ME2–CoA interaction as a critical regulator of muscle energy metabolism. Together, these findings position ME2-CoA as a mitochondrial redox sensor that converts oxidative stress signals into a protective metabolic response. Furthermore, they suggest that pharmacological targeting of the ME2 exo site may represent a therapeutic strategy for skeletal muscle disorders and other diseases characterized by mitochondrial ROS and NADPH imbalance.

A notable aspect of this study is the identification of CoA disulfide in mammalian cells. While CoA disulfide has long been recognized as a redox regulator in certain bacterial species ^23^, it has generally been assumed that eukaryotic systems lost this mechanism during evolution. Our findings challenge this view by demonstrating that CoA disulfide retains a functional role in mammalian mitochondria through its interaction with and regulation of ME2. This suggests a model that CoA disulfide cooperates with canonical eukaryotic redox systems, such as the glutathione (GSH/GSSG) pathway, to buffer mitochondrial oxidative stress. Whether CoA disulfide plays regulatory roles outside mitochondria remains an open question. Notably, ME2 has a cytosolic isoform, ME1, which exclusively utilizes NADP□ as a cofactor. However, ME1 was not identified as a CoA-binding candidate in our proteomic screen, suggesting that ME1 is likely not regulated by CoA. Consistent with this, the ME2 exo site CoA-binding residues are not conserved in ME1.

This work also provides the first direct evidence linking ME2 to exercise physiology. Prior studies have largely focused on ME2’s roles in cancer and neurological disorders.^27,30,36,37^ As a key metabolic enzyme, ME2 would be expected to function prominently in tissues with high energetic demand, such as skeletal muscle. Skeletal muscle cells also generate a lot of oxidative stress during exercise.^32,34^ Thus, skeletal muscle depends on ME2 function for optimal performance. Consistent with this notion, mice with *Me2 K197E* mutant, which lost the CoA-mediated redox regulation but retained the basal enzymatic activity, displayed minimal phenotypes under basal metabolic conditions, as assessed by metabolic cage analyses. In contrast, under metabolic stress induced by treadmill exercise, the functional consequences of disrupting the ME2 exo site became pronounced. These findings suggest that the regulation of ME2 by CoA functions primarily as a stress-responsive metabolic module rather than as a constitutive energy-producing pathway.

This stress-specific role is also consistent with observations that ME2 expression is elevated in certain cancers ^31^, which typically experience heightened oxidative stress compared with normal tissues ^38^. Despite ME2’s established role as an oncogenic factor in multiple cancer types, we did not observe a strong proliferative phenotype associated with the ME2 exo site mutation in cultured cancer cells. It is possible that standard cell culture conditions provide relatively low oxidative stress compared with the physiological tumor microenvironment, where oxygen and nutrient limitations exacerbate ROS production. These factors likely obscure the contribution of the ME2-CoA axis in vitro, highlighting the need for future in vivo studies to fully define its role in tumor progression.

Our work expands the functional repertoire of CoA beyond its traditional role as a metabolic cofactor and establishes it as a redox sensor and direct protein regulator in mitochondrial redox control. This function enables cells to sense both the availability of CoA and the level of oxidative stress to mount an adaptive and effective metabolic response to oxidative stress, particularly during energetically demanding conditions such as exercise. These findings identify the ME2–CoA axis as a critical component of mitochondrial redox homeostasis and open new avenues for therapeutic intervention in diseases driven by mitochondrial oxidative imbalance.

## Method

### Reagents

D-pantothenic acid hemicalcium salt (21210), p-anisaldehyde dimethyl acetal (10445), camphorsulfonic acid (C2107), 1-amino-3-butyne (715190) and azide-PEG3-biotin (762024) were purchased from Sigma-Aldrich. Human/mouse mitochondrial malic enzyme antibody (sc-12399) and Anti-Rabbit antibody conjugated with horseradish peroxidase (sc-7074) antibodies were purchased from Cell Signaling Technology. Protease inhibitor cocktail, □^12^C_6_, ^14^N_2_]-L-lysine, □^12^C_6_, ^14^N_4_]-L-arginine, □^13^C_6_, ^15^N_2_]-L-lysine and □^13^C_6_, ^15^N_4_]-L-arginine were purchased from Sigma-Aldrich. Dialyzed FBS, FBS, DMEM, SILAC grade DMEM and RPMI media were purchased from Thermo Scientific. Ham’s-F12 media was purchased from Corning. Sequencing grade modified trypsin was purchased from Promega. ECL plus western blotting detection reagent (80196) and disuccinimidyl suberate (21655) were purchased from Thermo Scientific. Sep-Pak C18 cartridge was purchased from Waters. Amicon Ultra centrifugal filter unit with ultracel-10 membrane was purchased from EMD Millipore. 2-2(pyridyl)ethyl silica gel (53798), Coenzyme A-reduced (C3019) and oxidized (C2643), L-malate (M9138) and fumarate (F1506) were purchased from Sigma-Aldrich.

### Biotin-CoA Synthesis

We first synthesized PMB-protected pantothenate ^39^. A flask with a stir bar and D-pantothenic acid hemicalcium salt (10 g, 42.0 mol) was flushed with nitrogen gas for 20 min. Anhydrous DMF (100 mL) was added followed by concentrated H_2_SO_4_ (1.3 mL dropwise, 42 mmol). The mixture was stirred for 15 min at room temperature. Then p-anisaldehyde dimethyl acetal (7.2 mL, 42 mmol) and camphorsulfonic acid (1 g, 4.6 mmol) were added and the reaction was stirred overnight. The reaction mixture was extracted at least three times with 75 mL ethyl-acetate and 100 mL water. The organic layers were combined and washed five times with 250 mL water, dried with Na_2_SO_4_ and evaporated. The resulting white solid was washed with 150 mL of cold (-20 □C) dichloromethane to remove any remaining p-anisaldehyde dimethyl acetal. The desired product was obtained as a white crystalline solid. 1H-NMR (CDCl_3_, 300 MHz): δ= 7.39 (d, 2H), 6.81 (d, 2H), 5.40 (s, 1H), 4.15 (s, 1H), 3.80 (s, 3H), 3.65-3.40 (m, 4H), 2.60 (t, 2H), 1.09 (s, 6H). ESI-MS □M^-^] calcd. 336.3645 for C21H28N2O5, found 336.48.

Then Alkyne-tagged pantetheine analogue was made ^40^. To a flask flushed with nitrogen gas, PMB-protected pantothenate (434 mg, 1.28 mmol), PyBOP (732 mg, 1.41 mmol), and 1-amino-3-butyne (88mg, 1.28 mmol) were dissolved in dry dichloromethane (50 mL). Diisopropylethylamine (677 μL, 3.84 mmol) was then added, and the reaction was stirred overnight. The mixture was evaporated, and the resultant oil was purified by column chromatography (2: 1 Hex:EtOAc). The product was obtained as an oil. 1H-NMR (300 MHz, CDCl_3_) δ= 7.40 (d, 2H), 7.10 (m, 1H), 6.95 (m, 2H), 6.25 (m, 1H), 5.45 (s, 1H), 4.10 (s, 1H), 3.78 (s, 3H), 3.67 (m, 2H), 3.63-3.47 (m, 2H), 3.39-3.25 (m, 2H), 2.46 (t, 2H), 2.38 (t, 2H), 1.96 (m, 1H), 1.12 (s, 6H). The obtained product (87 mg, 0.22 mmol) was dissolved in 1M HCl:THF, 1:1 (10 mL) and stirred for 1 hr or until the starting material was gone by TLC, at which point 1M NaOH was added to neutralize the solution. The solvent was evaporated, and the product was purified by column chromatography (EtOAc to 15% MeOH-EtOAc). The desired compound was obtained as an oil. 1H NMR (400 MHz, CDCl_3_) δ 7.36 (s, 1H), 6.22 (s, 1H), 4.10 (q, 1H), 3.98 (s, 1H), 3.60 – 3.53 (m, 2H), 3.47 (d, 3H), 3.38 (qd, 2H), 2.45 (t, 2H), 2.39 (td, 3H), 1.00 (s, 3H), 0.91 (s, 3H). ESI-MS □*M*+Na]^+^ calcd 293.1433 for C13H22N2O4, found 293.10.

An enzymatic method was used to make Biotin-CoA. The CoA biosynthesis enzymes, pantothenate kinase (CoaA), phosphopantetheine adenylyltransferase (CoaD) and dephospho-CoA kinase (CoaE), were purified from the *E. coli* ASKA strain library. Pantetheine analogue (5 mM in 30 HEPES pH 8.7 buffer) was added to a mixture of 100 mM HEPES pH 8.7, 18 mM MgCl_2_, 18 mM ATP, 15 μM CoaA, 15 μM CoaD and 15 μM CoaE at 37 oC for 1 hr. The reaction was quenched with 1 eq acetonitrile (ACN) followed by centrifugation and purified by high-performance liquid chromatography (Buffer A: H_2_O with 0.1% TFA, Buffer B: ACN with 0.1% TFA) to yield the CoA analogue, a white powder. ESI-MS □*M*+H]^+^ calcd 760.1504 for C23H36N7O16P3, found 760.05. Click-chemistry was initiated by the addition of 3 mM sodium ascorbate to 2.86 mM CoA analogue, 2.76 mM biotin azide, 2 mM TBTA, and 2 mM CuSO_4_ in 90 mM HEPES pH 8.7 at 37 □C for 1 hr. After centrifugation, the reaction mixture was purified by high-performance liquid chromatography (Buffer A: H_2_O with 0.1% TFA, Buffer B: ACN with 0.1% TFA) to yield the Biotin-CoA, a white powder. ESI-MS □M/2+H]^+^ calcd 603.0305, found 603.12.

### Stable isotope labeling with amino acids in cell culture (SILAC)

For proteomic experiments, the ‘Light’ and ‘Heavy’ human embryonic kidney (HEK) 293T cell lysates were prepared. ‘Light’ HEK 293T cells were maintained in SILAC grade DMEM media supplemented with 100 mg/L □^12^C_6_, ^14^N_2_]-L-lysine, 100 mg/L □^12^C_6_, ^14^N_4_]-L-arginine, and 10% dialyzed FBS. ‘Heavy’ HEK 293T cells were similarly maintained in SILAC grade DMEM media supplemented with 100 mg/L □^13^C_6_, ^15^N_2_]-L-lysine, 100 mg/L □^13^C_6_, ^15^N_4_]-L-arginine, and 10% dialyzed FBS. Cells were cultured in the SILAC media for eight doubling times to achieve maximum incorporation of ‘labeled’ amino acids into proteins before preparing the lysate for the proteomics experiment. After eight passages, the cells were collected and lysed using lysis buffer comprising of 50 mM HEPES pH 7.4, 150 mM NaCl, 1% NP-40, 10% glycerol, and 31 1% v/v protease inhibitor cocktail. After quantifying protein concentration by Bradford assay, two tubes with 3 mg ‘heavy’ lysate were incubated with 800 μM Biotin-CoA and 3 mg ‘light’ lysate were incubated with either: (i) 800 μM biotin-azide or (ii) 800 μM biotin-CoA + 8 mM CoASH. After 30 min incubation, streptavidin beads were added and incubated for 1 hr. Supernatant was removed, and the beads were washed twice with washing buffer comprising of 50 mM HEPES pH 7.4, 150 mM NaCl, 0.2% NP-40, 10% glycerol. Each pair of samples (heavy labeled and light labeled) were mixed and then washed two more times. After washing, disulfide reduction and protein denaturation were performed in 6 M urea, 10 mM DTT, 50 mM Tris-HCl pH 8.0 at room temperature for 1 hr. Sulfhydryl alkylation was done by the addition of 40 mM iodoacetamide and incubation at room temperature for 1 hr. Alkylation was stopped with the addition of 3.8 mM DTT and incubation at room temperature for 1 hr. Samples were then diluted 7 times with 50 mM Tris-HCl pH 8.0, 1 mM CaCl_2_. Trypsin digestion was initiated by the addition of 30 μg trypsin and digestion was performed at 37 °C for 18 hr. Trypsin digestion was quenched with 0.2 % trifluoroacetic acid and the samples were desalted using Sep-Pak C18 cartridge and then lyophilized. The lyophilized peptides were dissolved in 2% ACN with 0.5% formic acid (FA) and nanoLC-MS/MS analysis was carried out on an LTQ-Orbitrap Elite mass spectrometer. Samples were loaded onto an Acclaim PepMap nano-Viper C18 trap column (5 μm, 100 μm x 2 cm, Thermo Dionex) for on-line desalting then separated on a C18 RP nano-column (5 μm, 75 μm x 50 cm, Magic C18, Bruker) at a flow rate of 0.3 μL/min. The HPLC gradient was 5-38% ACN with 0.1% FA for 120 min. The Orbitrap was operated in positive mode with spray voltage 1.6 kV and source temperature 275 °C. Data-dependent 32 acquisition mode was used by one precursor ions MS survey scan from m/z 300 to 1800 at resolution 60,000 using FT mass analyzer, followed by up to 10 MS/MS scans at resolution 15,000 on 10 most intensive peaks. All data were acquired in Xcalibur 2.2 operation software package.

### Validation of SILAC results

HEK 293T cells were lysed (50 mM HEPES pH 7.4, 150 mM NaCl, 1% NP-40, 10% glycerol) and 500 μg of protein lysate was aliquoted into 4 tubes. Three of the tubes were incubated with 500 μM biotin-CoA for 15 min, then different concentrations of CoASH (0 mM, 0.25 mM, 2.5 mM) were added. Streptavidin beads were added to all four tubes and the tubes were incubated at room temperature for 30 min. Beads were washed (50 mM HEPES pH 7.4, 150 mM NaCl, 0.2% NP-40, 10% glycerol) at least four times, and analysis by anti-ME2 immunoblotting was performed.

### Western Blot Analysis

Protein samples were separated by SDS-PAGE and transferred to PVDF membranes. The membranes were blocked using 5% BSA in PBST (25 mM Tris-HCl, pH 7.4, 150 mM NaCl, 0.1% Tween-20) and incubated with antibodies in 5% BSA in PBST. Western blotting was detected using ECL Plus reagents and Typhon Imager.

### Cell Culture

HEK 293T cells were cultured in complete DMEM medium supplemented with 10% heat-inactivated FBS. A549 cells were cultured in either RPMI or Ham-F12 media, both supplemented with 10% heat-inactivated FBS.

### SAXS experiments

Small-angle X-ray scattering (SAXS) data were performed on the beamline BL19U2 at the SSRF. Briefly, proteins were subjected to size exclusion chromatography in a buffer containing 20 mM Tris-HCl pH 7.4, 150 mM KCl, 1 mM DTT. 60 μl 1-2 mg/mL protein was used and the data were collected at 1.03 Å. All measurements were carried out in vacuum with exposure times of 1 s with 20 frames of each protein sample as well as buffer-only samples. All 2D scattering images were converted to 1D SAXS curves using the software package BioXTAS RAW^41^. The ATSAS package ^42^ was used for subsequent data processing and FoXS was used for atomistic modelling^43^. Due to the considerable experimental noise at higher scattering angles, only the most informative part of scattering curves between 0.01 Å^-1^ and 0.3 Å^-1^ was used for structural analysis.

### Differential scanning calorimetry (DSC)

DSC experiments were performed with a DSC calorimeter from MicroCal (Northampton, MA, USA) at a scan rate of 1.0□ min^-1^. Buffer (20mM Tris-HCl pH 7.4, 30 mM KCl) was used in the reference cell of the calorimeter. Proteins were dialyzed on the buffer and used at a concentration of 1 mg/ml. Data were analyzed with the MicroCal DSC standard analysis software.

### Construction of ME2 Plasmids

To make the cDNA library, total RNA was isolated from HEK 293T using the RNeasy 33 mini kit (Qiagen, 74106) according to the manufacturer’s instructions. The isolated total RNA was used for the cDNA synthesis using RT-PCR. Flag-tagged ME2 (19 – 584) was PCR-amplified from HEK 293T cDNA using: taattaatcatatgttgcacataaaagaaaaaggcaagccacttatgc (forward primer) and taattaatctcgagctaCTTATCGTCGTCATCCTTGTAATCttctgttatcacaggagggcttgatg (reverse primer with Flag-tag). ME2 was cloned into *Nde* I/*Xho* I restriction sites of pET28a and was sequence-confirmed. Arginine-to-glutamate ME2 mutant vectors were constructed using a quick-change site-directed mutagenesis strategy with the following primers: R197E ME2: ggaatacggcctgatGAGtgcctgccagtgtgt (forward primer), atcaggccgtattcctgcacaagctgtatacaa (reverse primer). R556E ME2: aaatatgttaaagaaGAGacatggcggagtgaa (forward primer), ttctttaacatatttggccttgtcttcaggttc (reverse primer).

### Expression and purification of WT, R197E, and R197E/R556E ME2 in *E.* coli for biochemical assays

The sequence-confirmed pET28a plasmid was overexpressed in SoluBL-21. Cells were cultured at 37 □C in LB media with 50 μg/mL kanamycin. ME2 was purified using Ni-NTA affinity followed by size-exclusion chromatography. ME2 expression was induced by adding 200 μM isopropyl β-D-1-thiogalactopyranoside and cells were further incubated for 3 hr. Cells were collected at 11,000 g for 7 min and the cell pellet was stored at -80 □C until use. Cells were thawed and suspended in 50 mM HEPES pH 7.4, 500 mM NaCl, 30 mM imidazole, 1 mM MgCl_2_, 1 mM PMSF. Cells were lysed using an EmulsiFlex-C3 cell disruptor and then centrifuged at 48,384 *g* for 45 min using a Beckman Coulter refrigerated floor centrifuge. The soluble fraction was loaded onto a Ni-NTA agarose column then washed with 50 mM HEPES pH 7.4, 500 mM NaCl, 30 mM imidazole, 1 mM MgCl_2_. ME2 was eluted using 500 mM imidazole in wash buffer and concentrated for size-exclusion chromatography, which was performed using an ÄKTA pure chromatography system and a buffer of 50 mM HEPES pH 7.4, 500 mM NaCl, 1 mM MgCl_2_. Fractions containing pure ME2 (determined by SDS/PAGE analysis) were collected, concentrated, and stored at −80 °C. All mutants of ME2 were purified similarly.

### Thermal Shift Assay

In PCR tubes, 10 μM WT ME2 was mixed with or without 0.5 mM CoASH in a buffer of 50 mM HEPES pH 7.4, 500 mM NaCl, 1 mM MgCl_2_. The tubes were then put in a PCR thermocycler with the temperature oscillating from 32 to 52 °C for different tubes. After incubation for 10 min, the samples were then transferred to 1.5 mL tubes and centrifuged at 17,000 g for 15 min to remove denatured proteins. The supernatant was analyzed by 12% SDS-PAGE and proteins were visualized by Coomassie blue staining.

### Biotin-CoA pulldown of Recombinant ME2 Proteins

WT, R197E, and R197E/R556E ME2 at 2 μM were mixed with 50 μM Biotin-CoA in a buffer containing 50 mM HEPES pH 7.4, 500 mM NaCl, 1 mM MgCl_2_ for 15 min and then incubated with streptavidin beads for 30 min. Beads were washed at least three times with buffer and then boiled in 95 °C and analyzed by 12% SDS-PAGE and Coomassie blue staining.

### ME2 Activity Assay

Enzyme activity was measured by monitoring the formation of NADH at 340 nm using a UV-visible spectrophotometer (Cary 50 UV-visible spectrophotometer or BioTek Cytation 5). The reaction mixture contained 35 mM HEPES 7.4, 130 mM NaCl, 20 mM MgCl_2_, 10 mM KCl, and 50 nM ME2. The mixture was incubated for 4 min at 37 □C in the presence or absence of a CoA molecule or fumarate (4 mM), then NAD was added, and the reaction was initiated by adding L-malate. All assays were done at 37 □C. For ME2 enzyme activity under different redox conditions (GSH/GSSG), either CoA or CoA disulfide was mixed with GSSG or GSH and incubated with ME2 enzyme at 37 □C for 10 mins. For relatively specific 36 activity assays, 1 mM NAD and 1 mM L-malate were used. CoASH and all acyl-CoAs were assayed at 100 μM. CoA-S-S-CoA was tested at various concentrations with maximum activation observed at 10 μM. The kinetic parameters were determined by using 10 mM L-malate with 0.05-1 mM NAD and 1.5 mM NAD with 0.5-20 mM L-malate. Data was analyzed using GraphPad Prism software.

### Protein Cross-Linking

WT or R197E ME2 at 20 μM was incubated with 10 μM CoA-S-S-CoA in a buffer containing 50 mM HEPES pH 7.4, 500 mM NaCl, 1 mM MgCl_2_ for 15 min. Crosslinking was done by the addition of 0.5 mM or 1 mM disuccinimidyl suberate (DSS) and incubation at room temperature for 30 min. The crosslinking reactions were stopped with the addition of 50 mM Tris pH 8.0 and incubation at room temperature for 15 min. Samples were analyzed by 8% SDS-PAGE and visualized by Coomassie blue staining.

### Cloning, expression and purification of ME2 for crystallization

ME2 (without the mitochondrial signal peptide 1-18) and mutants were overexpressed in *Escherichia coli* BL21 (DE3) cells (Novagen) using a modified pET 28a-TEV vector with a T7 promoter system. Cells were grown in LB at 37°C until OD600 reached 0.6 and then induced for 12 h at 18°C with 0.2 mM isopropyl β-D-1-thiogalactopyranoside. The cultures were harvested by centrifuging at 4,000 g for 10 min at 4°C and then resuspended in lysis buffer (30 mM Tris-HCl pH 7.3, 1000 mM KCl and 50 mM imidazole) and disrupted by sonication. The lysate was centrifuged at 47,000 g for 35 min to remove cell debris. The soluble fraction was applied to a Ni^2+^-chelating column (GE Healthcare). After sample loading, the column was washed with washing buffer (30 mM Tris-HCl pH 7.3, 600 mM KCl and 50 mM imidazole) and then eluted with 5 column volume elution buffer (30 mM Tris-HCl pH 7.3, 300 mM KCl and 200 mM imidazole). The His-Tag was cleaved using His-TEV protease at 4°C for 8 h and removed by a Ni^2+^-chelating column. Further purification was performed by size-exclusion chromatography using a Superdex 200 (GE Healthcare) with the final buffer (30 mM Tris-HCl pH 7.3, 150 mM KCl). Peak fractions were collected and analyzed by SDS-PAGE. All purified proteins were centrifuged to 8 mg/ml for crystallization.

### Crystallization, data collection and structure determination

About 7 mg/mL ME2 protein was in a solution containing 30 mM Tris-HCl pH 7.3, 150 mM KCl and 5% glycerol. Crystals of apo-ME2 were obtained against a reservoir solution containing 25% PEG 3350, 0.1 M HEPES pH 7.5, 0.2 M lithium sulfate with the hanging-drop vapour-diffusion method. To obtain the complex crystals of ME2 with CoASH or CoA disulfide (CoASSCoA), 1 mM final concentration of CoASH or CoA disulfide was added to the purified ME2 protein. The CoASH and CoA disulfide complexes were crystallized against reservoir solutions composed of 25% PEG 3350, 0.1 M HEPES pH 7.0, 0.2 M lithium sulfate and 25% PEG 3350, 0.1 M Tris-HCl pH 8.5, 0.2 M ammonium acetate, respectively. For the three quinary complex with CoASH/NAD^+^/PYR/Mn^2+^ or CoA disulfide/NAD^+^/PYR/Mn^2+^, ME2 or K156T mutant protein was concentrated and buffer exchanged to 20 mM Tris-HCl pH 7.4, 30 mM KCl, 1 mM DTT, 10 mM NAD^+^, 5 mM MnCl_2_, 10 mM pyruvate and 1-10 mM CoASH/CoA disulfide. The two quinary ME2 WT complexes were crystallized against a reservoir solution containing 20% PEG 3350, 0.2 M ammonium tartrate pH 7.0 and 14% PEG 3350, 0.1 M Bis-Tris pH 7.0, 0.2 M sodium formate, respectively and the ME2 K156T-CoA disulfide/NAD^+^/PYR/Mn^2+^ structure was crystallized against reservoir solutions composed of 12% PEG 3350, 0.1 M sodium malonate, pH 7.0 (final pH 7.4).

Crystals were flash cooled in liquid nitrogen in relative mother liquor containing 25% glycerol as a cryoprotectant. Data were collected on beamlines BL17U1 and BL18U1 at Synchrotron Radiation Facility (SSRF) and processed by HKL2000 ^44^. Structures of apo-ME2 and ME2/cofactor/ substrates complexes were solved by the molecular replacement protocol with the program PHASER. Data collection and processing statistics were shown in supplementary Table 2.^45^

### Generation of ME2 Stable KD Cell Lines

*ME2* shRNA lentiviral plasmids in pLKO.1-puro vector were purchased from Sigma-Aldrich. *ME2* shRNA (TRCN0000294007, 3UTR region): CGGAGTTCTTACAGAGCTACTAAACTCGAGTTTAGTAGCTCTGTAAGAAC TTTTTTG was used. After co-transfection of *ME2* shRNA plasmid, pCMV-ΔR8.2, and pMD2.G into HEK 293T cells, the medium was collected to infect A549 cells.

The ME2 KD cells were selected using 2 μg/mL puromycin in RPMI or Ham’s-F12 media supplemented with 10% heat-inactivated FBS. Cells infected with lentivirus containing control shRNA plasmid were carried out similarly ^46^.

### Confocal microscropy

*ME2* WT or knockdown cells were seeded in a 35mm dish and tranfected either ME2 WT or R197E mutant. After 24 hrs, cells were covered with 1μM MitoSOX (Thermo cat# M36006) solution in HBSS and incubated for 30 mins at 37°C and 5% CO2. Cells were washed three times with HBSS before covered with DAPI Fluoromount-G. The images were acquired using Leica stellaris 8 with 63x oil lense.

### Extraction of CoA molecules from mammalian cells

All the extraction solutions, buffers and solvents used for the CoA extraction were pre-cooled on ice. Cells or mitochondria were spiked with 0.2 nmol of d3-acetyl-CoA as an internal standard and were homogenized with 0.5 mL of methanol/water (1:1) containing 5% acetic acid (extraction buffer) using a Dounce homogenizer (30 strokes) on ice. The cell homogenates were centrifuged at 17,000 g for 15 min at 4 °C. The clear supernatant was loaded on a 3-mL ion exchange cartridge packed with 100 mg of 2-2(pyridyl)ethyl silica gel. The cartridge had been pre-activated with 3 mL of methanol and then with 3 mL of extraction buffer. The ion exchange resin was washed with 2 mL of extraction buffer to remove unbound metabolites. The acyl-CoAs trapped on the silica gel cartridge were eluted twice with 0.5 mL of methanol/250 mM ammonium formate (4:1). The combined effluent was lyophilized and stored at −80 °C until LC-MS analysis.

### LC-MS analysis of extracted CoA molecules

The lyophilized CoA samples were re-suspended in 150 μL of 50 mM ammonium acetate pH 6.8, centrifuged at 17,000 g for 5 min and 10 μL were injected. An 38 UltiMate3000 RSLC (Thermo) was coupled to an OrbitrapElite mass spectrometer for metabolite separation and detection. A XTerraMS C18 column (150 mm x 2.1 mm i.d., 3.5 μm, 155 Å) (Waters) was used with a flow rate of 200 μL/min. The solvent A was water with 5 mM ammonium acetate pH 6.8 and solvent B was methanol. The solvent gradient was as follows: 0 min, 2% solvent B; 2 min, 2% solvent B; 4 min, 15% solvent B; 7.5 min, 95% solvent B; 17.5 min, 95% solvent B; 18 min, 2% solvent B; and 24 min, 2% solvent B. The OrbitrapElite operated in ESI negative ion FT mode with Ion Max-S API source under Xcalibur2.2. The following parameters were used: ESI voltage -4.0 kV, Sheath gas: 40 (arbitrary unit), Aux gas: 0; Sweep gas: 0, Source temperature: 325 °C, S-Lens RF level: 50%. A full scan in FTMS was set at 120,000 resolving power across 300 to 1600 (m/z) in profile mode followed by two MS/MS DDA scans acquired from 0 min to 24 min. All data were analyzed using Xcalibur2.2 software package.

### HPLC detection of CoASH and CoA disulfide

Free CoASH was incubated with or without the addition of ME2 enzyme at 37 °C for 30 mins. The reaction mixture was then heated at 65 °C for 15 mins to denature the enzymes. After centrifuging at max speed for 10 mins, the supernatant was mixed with 1% FA at 1:1 ratio prior to injection into the HPLC system (Shimazu). The same gradient as the LC-MS method described above was used and UV absorbance at 260nm was used to detect the CoASH and CoA disulfide.

### Generation of *Me2 K197E* knock-in mice

The *Me2 K197E* knock-in mice were generated with a C57BL/6 background by CRISPR/Cas9-mediated genome editing at the Cornell University Stem Cell and Transgenic Mouse Facility. The targeted AAG-to-GAG substitution (encoding a Lys197Glu mutation, K197E) was confirmed by PCR and Sanger sequencing. The sequences of genotyping primers are: Forward 5’-GCTGTTGTGGTGACTGATGG-3’; Reverse 5’-GGAGAGATGCACTCACCATGT-3’. Mice were maintained under specific-pathogen-free (SPF) conditions in a barrier facility and housed in individually ventilated cages with ad libitum access to standard chow. All procedures involving animals were approved by the Institutional Animal Care and Use Committee.

### Mice treadmill test^47^

Adult wild-type (WT) and *Me2 K197E knock-in* mice (8–15 weeks of age) were subjected to treadmill training and endurance testing. Mice were first acclimated to the treadmill over three consecutive days. During each training session, animals were run at gradually increasing speeds, beginning at 8 m/min and increasing to 10 m/min on day 1, and to 12 m/min on days 2–3. Each speed interval was maintained for 5–10 minutes, with total daily training durations of approximately 15–30 minutes per mouse. Following 3 days of training, mice underwent a standardized treadmill endurance test. The protocol consisted of stepwise increases in running speed as follows: 12 m/min for 1 minute, 16 m/min for 1 minute, 18 m/min for 5 minutes, 20 m/min for 25 minutes, 22 m/min for 15 minutes, 24 m/min for 15 minutes, and 26 m/min for 13 minutes. Exhaustion was defined as the inability to continue running, indicated by remaining in the fatigue zone for more than 5 seconds. Time to exhaustion and total distance traveled were recorded. The endurance test was repeated for three consecutive days for each mouse. The blood serum and muscle samples were collected at the end of treadmill test of day 2 and 3 respectively.

### Metabolic cage

Mouse experiments were performed according to procedures approved by IACUC. Adult male wild-type (WT) and *Me2 K197E knock-in* mice (8–15 weeks of age) were housed at 22□°C under a 12-h light/dark cycle and given free access to food and water. All mice used for indirect calorimetry experiments (n=8/genotype) were housed in groups at 22□°C with bedding and shredded paper strips in the cage until experimental intervention. Mice (12 weeks of age) were then placed, single housed, in metabolic cages (Sable Systems International, Promethion high-definition behavioural phenotyping system) housed in thermal cabinets set to 22□°C with a 12-h light/12-h dark schedule (light 07:00 to 19:00). Mice had ad libitum access to diet and water and were allowed to acclimatize in the metabolic cages for 3□d. Energy expenditure (EE), respiratory exchange ratio (RER), locomotor activity, food/water consumption and total body mass were recorded every hour using Sable Systems data acquisition software (IM-3 v.20.0.3). At hour 90, mice underwent a cold challenge at 6°C until the end of the experiment, resulting in increased energy expenditure. Statistical analysis was performed using two way ANOVA with Tukey’s multiple comparisons test.

## Author Contribution

W.C., O.D.N., X.R.L., and M.F.Z. contributed equally to this work. W.C., O.D.N., and H.L. conceived the project and designed the research. O.D.N. performed the SILAC experiments. W.C. and O.D.N. performed the in vitro enzyme reactions; W.C. performed the cellular studies and confocal imaging. X.R.L. and M.F.Z. performed small-angle X-ray scattering experiments, conducted structural analysis, and drafted the structural sections of the manuscript. X.L., T.Y., S.Z., and C.H.Y. generated the CRISPR knock-in mice. W.C., X.L., A.F., and B.H.A. performed the mouse experiments. M.S., E.L. and J.F.R. conducted the metabolic cage experiments. Y.G.Z. and O.D.N. performed the cellular cancer experiments. C.H.Y. performed the metabolomics analysis and data interpretation. W.C., O.D.N., Z.Z.C., and H.L. wrote the manuscript with input from all authors. Z.Z.C. and H.L. provided supervision and funding.

## Acknowledgement

This work is supported in part by funding from Howard Huges Medical Institute, National Institutes of Health/National Institute of General Medical Sciences (R35GM131808), and University of Chicago Comprehensive Cancer Center. O.D.N was supported by National Institutes of Health/National Institute of General Medical Sciences grant 5T32GM008500. Z.C. is supported by National Key R&D Program of China (2023YFF1206001) and National Natural Science Foundation of China (32371268).

## Data Availability

X-ray crystal structure data are deposited in Protein Daba Bank with the following PDB ID: 25EH, 25EI, 25EJ, 25EK, 25EL.

## Supplementary Figures

**Figure S1.**
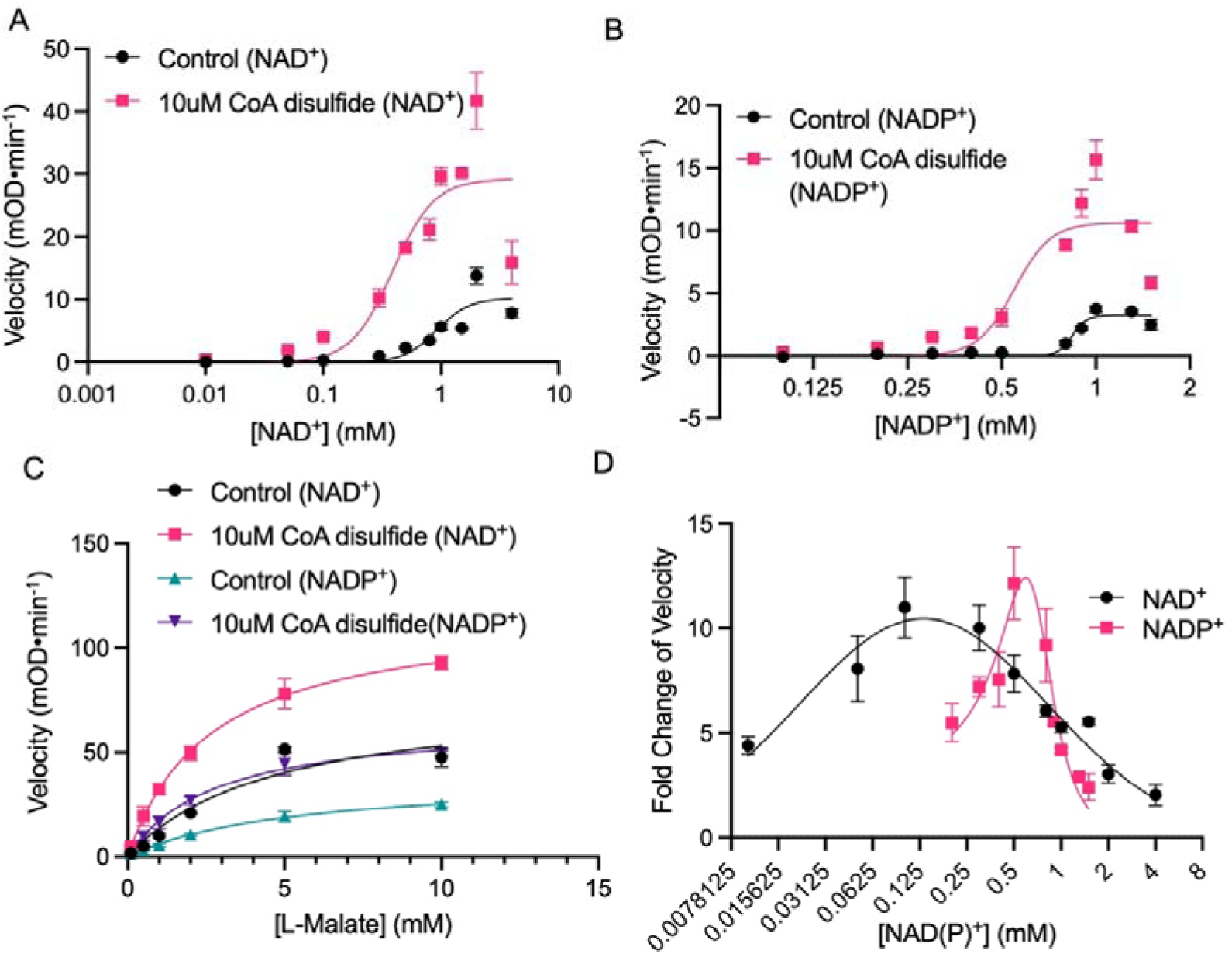
ME2 enzyme activity under various substrate concentrations. ME2 enzymatic activity with and without CoA disulfide was measured at various (A) NAD^+^, (B) NADP^+^, (C) L-malate concentrations while keeping other substrate concentration constant. (D) CoA disulfide promotes both NAD^+^ and NADP^+^-dependent ME2 enzyme activity, but the degree of activation varies at different nucleotide concentrations.

**Figure S2.**
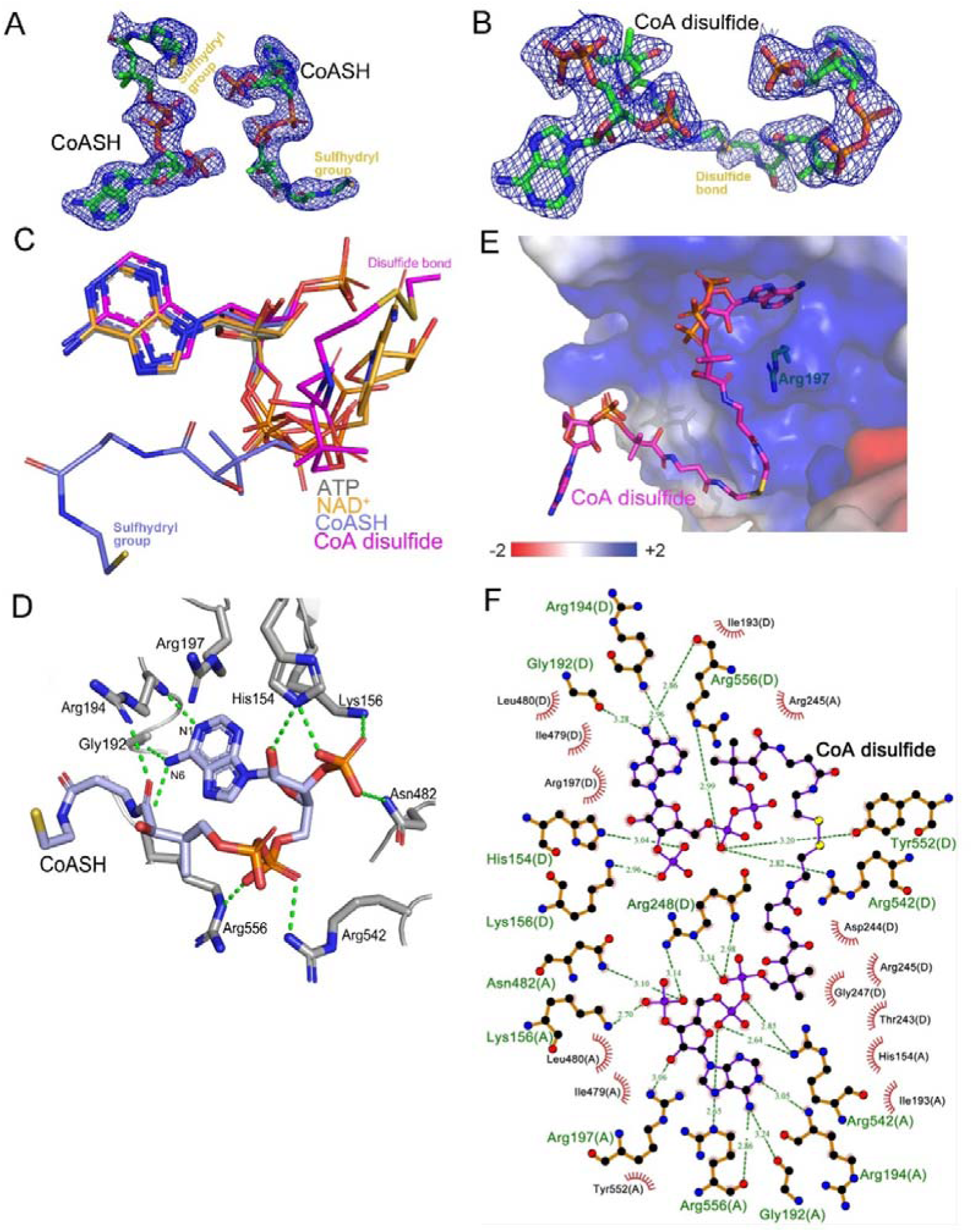
Structural analysis of CoASH and CoA disulfide binding to ME2 and ligand comparison at the exo site. *2Fo-Fc* electron density maps for the two CoASH molecules of ME2-CoASH complex structure (A) and a CoA disulfide molecule of ME2-CoA disulfide complex structure (B). The electron density is displayed as a blue mesh (contoured at 1.0 σ). (C) Schematic drawing showing the overlay of the ATP (gray), NAD^+^ (orange), CoASH (slate) and CoA disulfide (magenta) bound at the exo site of ME2. (D) The binding site of CoASH in the ME2-CoASH complex structure. CoASH (slate) and interacting protein residues (gray) are shown as sticks. Hydrogen bonds between the protein and the cofactor are indicated as green dotted lines. (E) The surface potential of the ME2-CoA disulfide complex generated by the APBS tool of PyMOL (unit: kT/e) was displayed as a color gradient ranging from red (negative) to blue (positive). Detailed interactions and hydrogen bonds between ME2 and CoA disulfide in the ME2-CoA disulfide binary complex structure (F) are shown as the LigPlot diagram, note that different chains were labelled in the parentheses. The green dashed lines correspond to the hydrogen bonds. Spoked arcs represent hydrophobic contacts.

**Figure S3.**
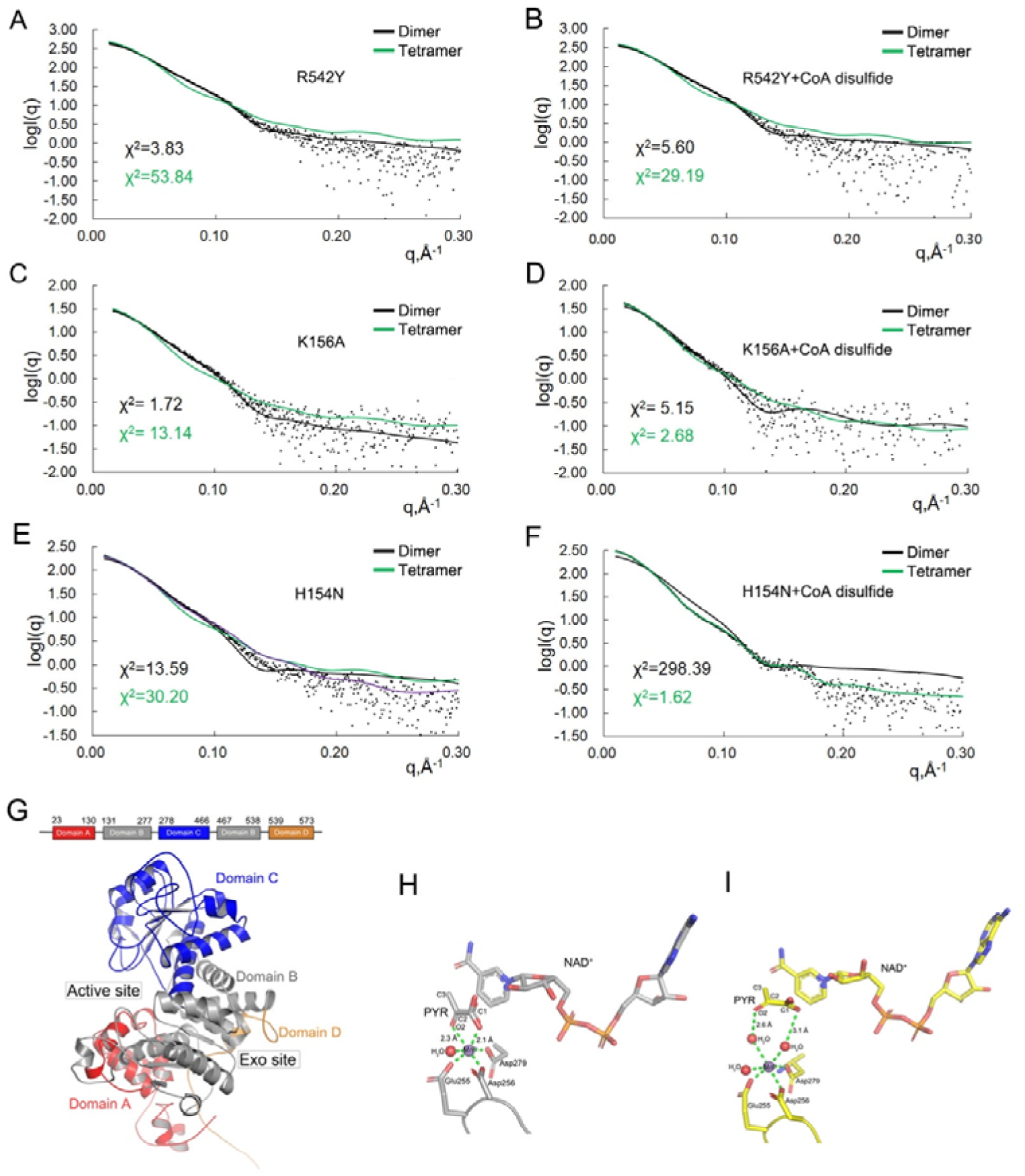
SAXS analysis of ME2 mutants reveals oligomeric state in the presence of CoA disulfide and structural comparison with ME2-CoASH complexes. Overlay of experimental scattering profiles with calculated scattering profiles for R542Y (A, B), K156A (C, D) and H154N (E, F) mutants, alone and in the presence of CoA disulfide. Experimental data are represented in black dots. The theoretical scattering curves of dimers (black lines) and tetramers (green lines) are shown. (G) Schematic domain and a cartoon representation of the apo-ME2 protomer. The domains A, B, C and D of the structure are shown in red, gray, blue and brown, respectively. The active site and exo site of the protomer are also indicated. Schematic drawing of the active site in the crystal structure of ME2 in complex with CoA disulfide/NAD^+^/PYR/Mn^2+^ (H) and CoASH/NAD^+^/PYR/Mn^2+^ (I). The manganese ion and the ligated water are shown as spheres.

**Figure S4.**
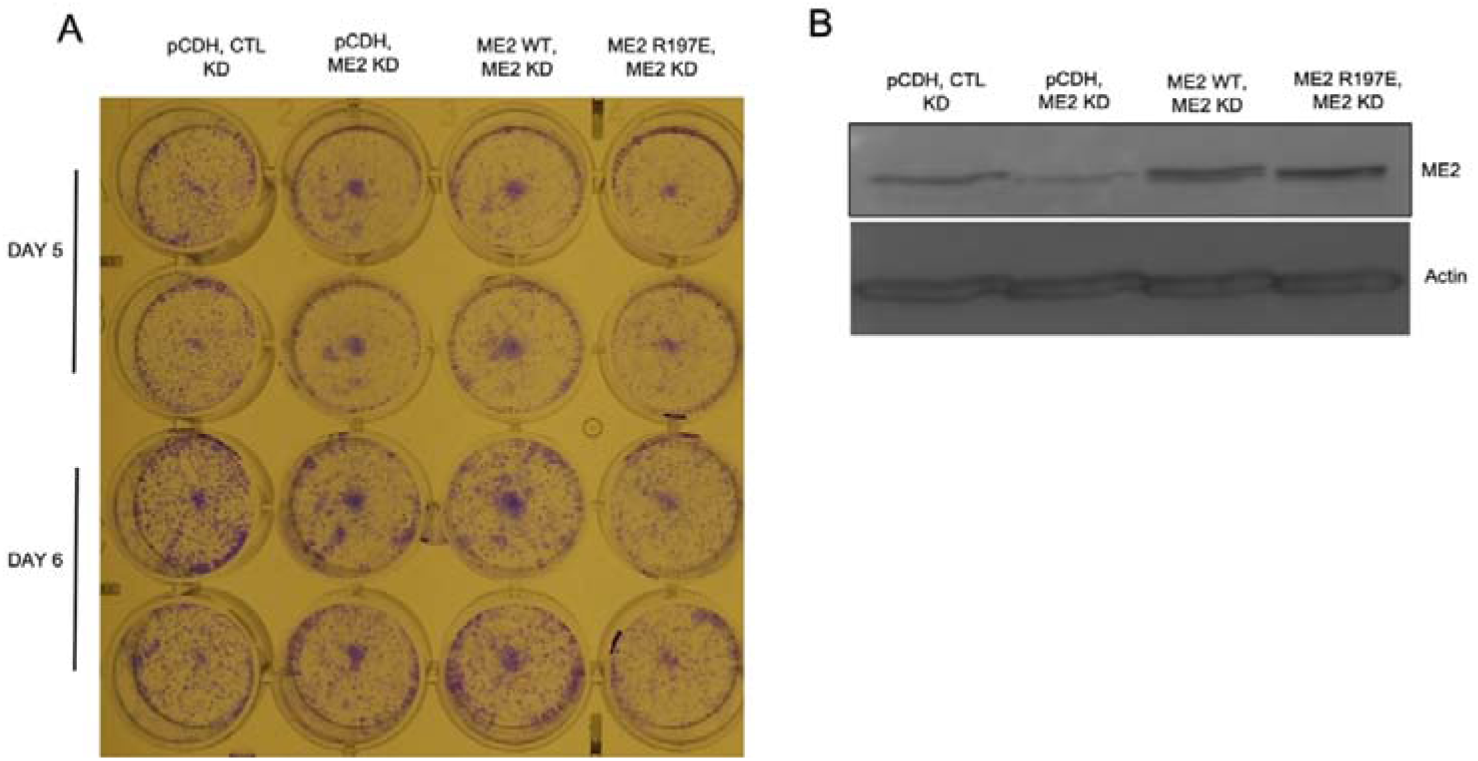
The effect of ME2 R197E on A549 cells colony formation ability. A549 control knockdown and ME2 knockdown cells were stably transfected with pCDH, ME2 WT, or ME2 R197E as indicated. Colony formation assay was performed to measure the tumorigenesis effect on ME2 knockdown and ME2 WT or R197E re-expression. (A) Representative images of colony formation assay at day 5 and 6; (B) Western blot for the validation of ME2 knockdown and ME2 WT or R197E re-expression.

**Figure S5.**
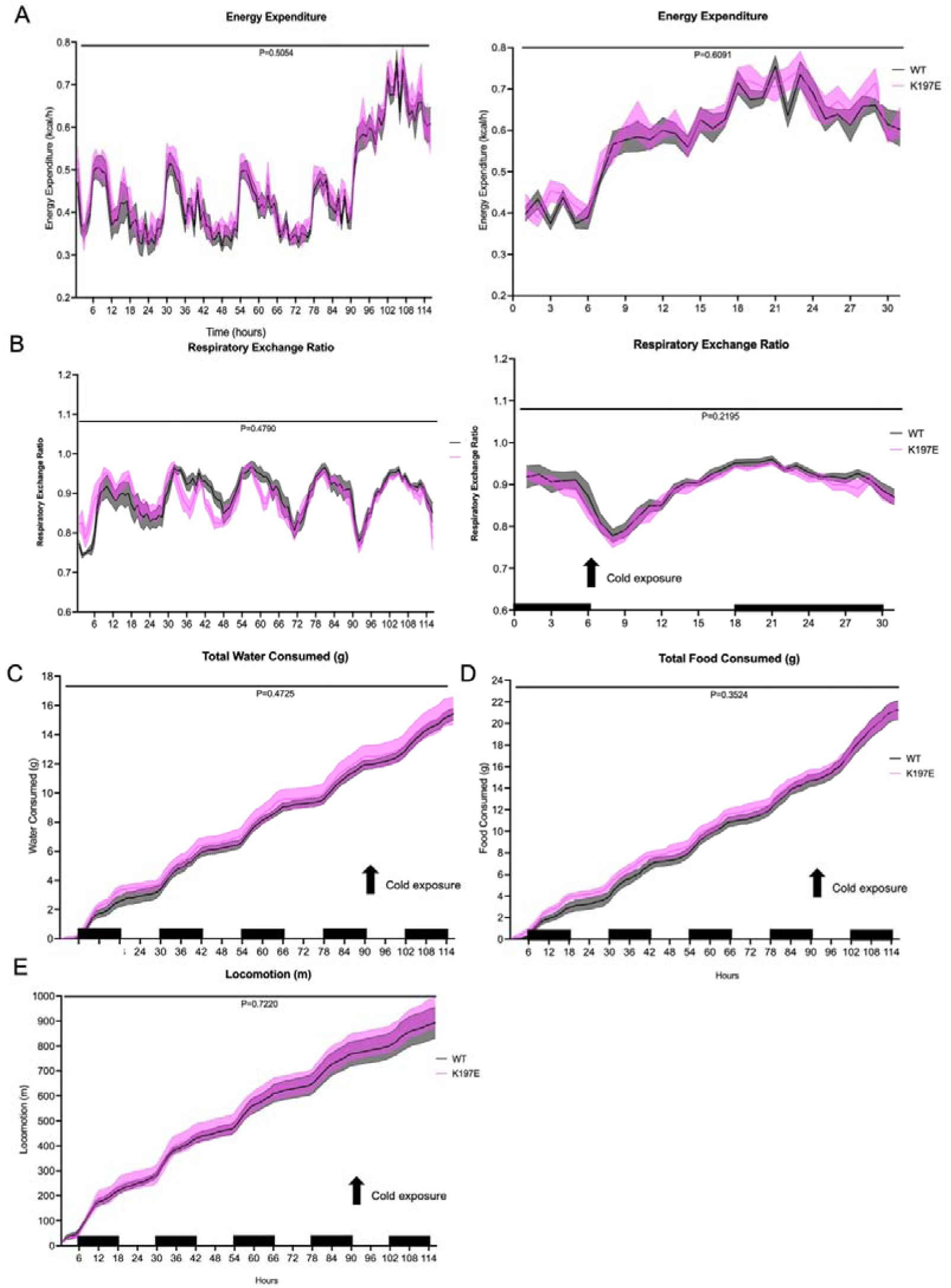
Metabolic cage result for *Me2 WT* (gray) and *K197E* mice (purple). The *Me2 K197E* CRISPR knock-in mice were generated on a C57BL/6J background and subject to a metabolic cage experiment. (A) Energy expenditure, (B) Respiratory exchange ratio, (C) Total water consumption, (D) Total food consumption and (E) Locomotion were monitored for 114 hours.

**Supplementary Table 1.**
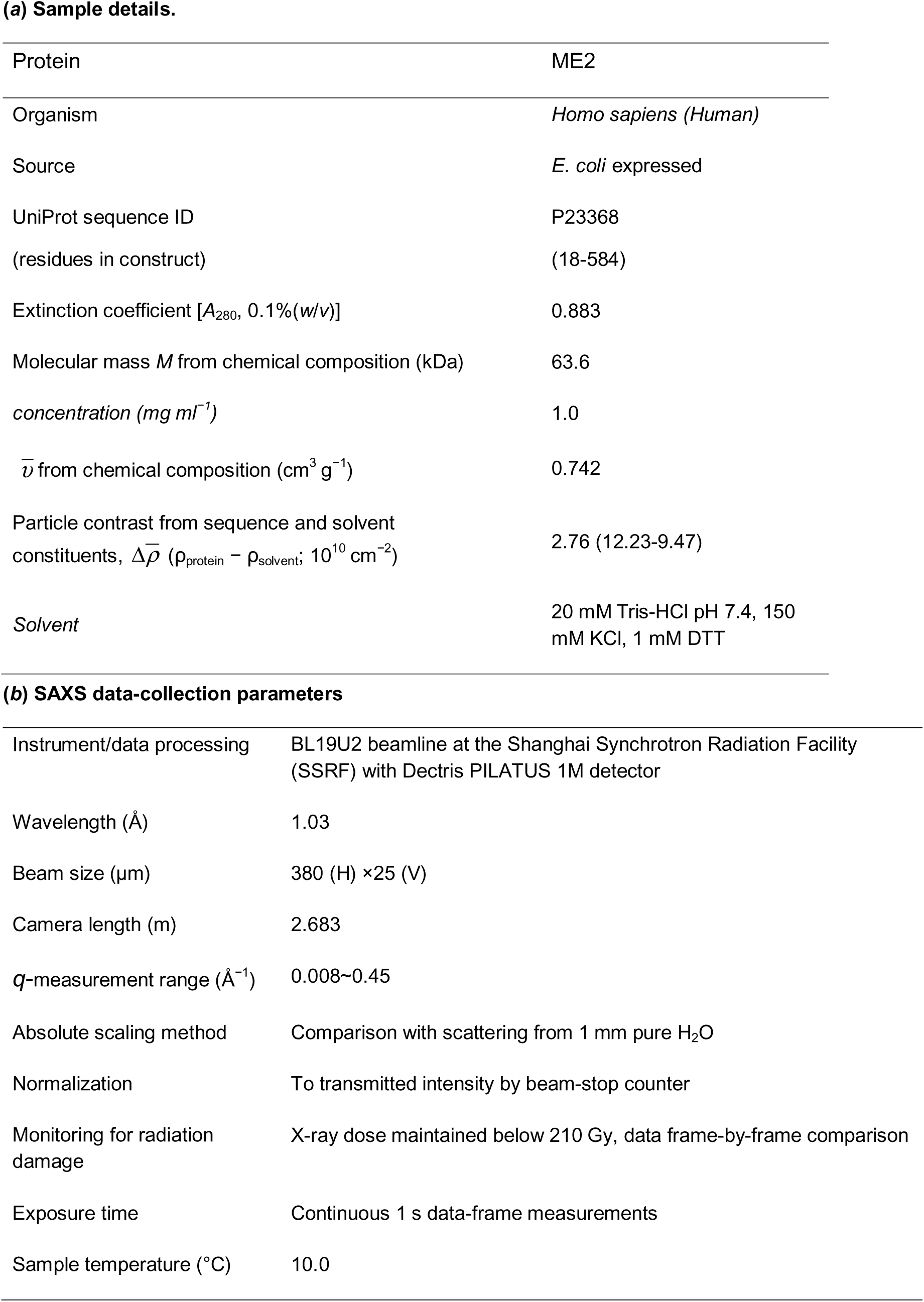
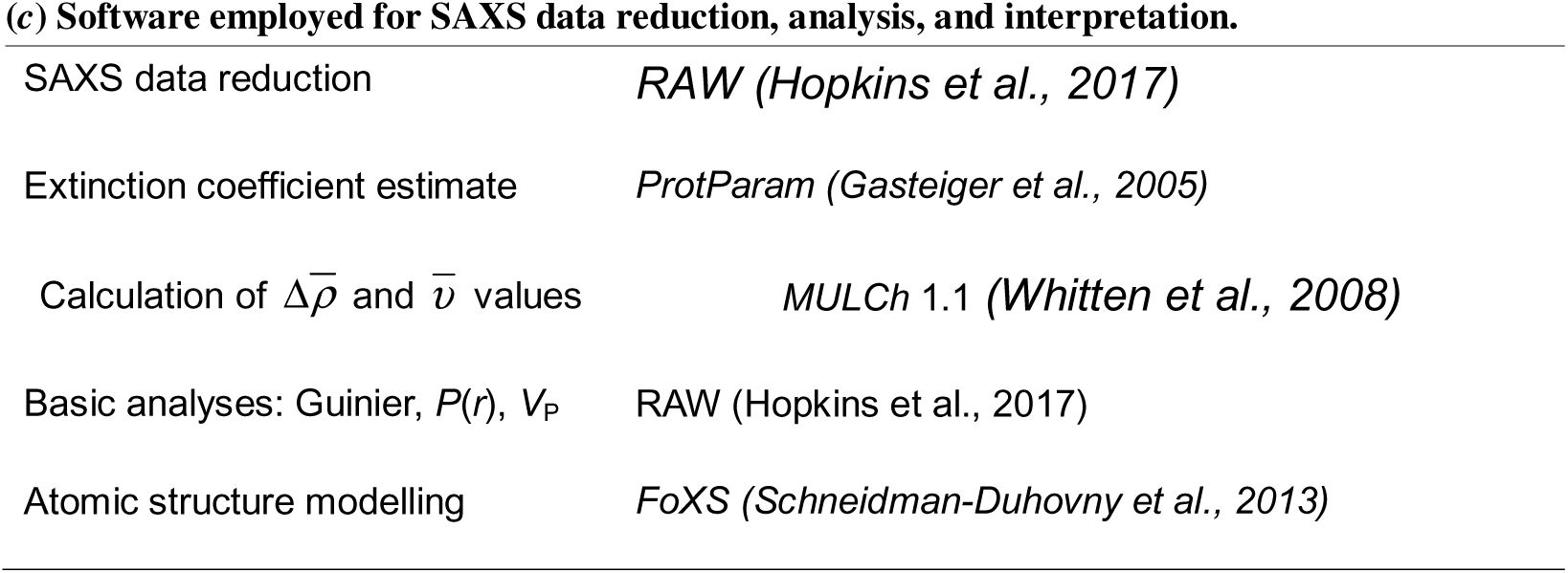
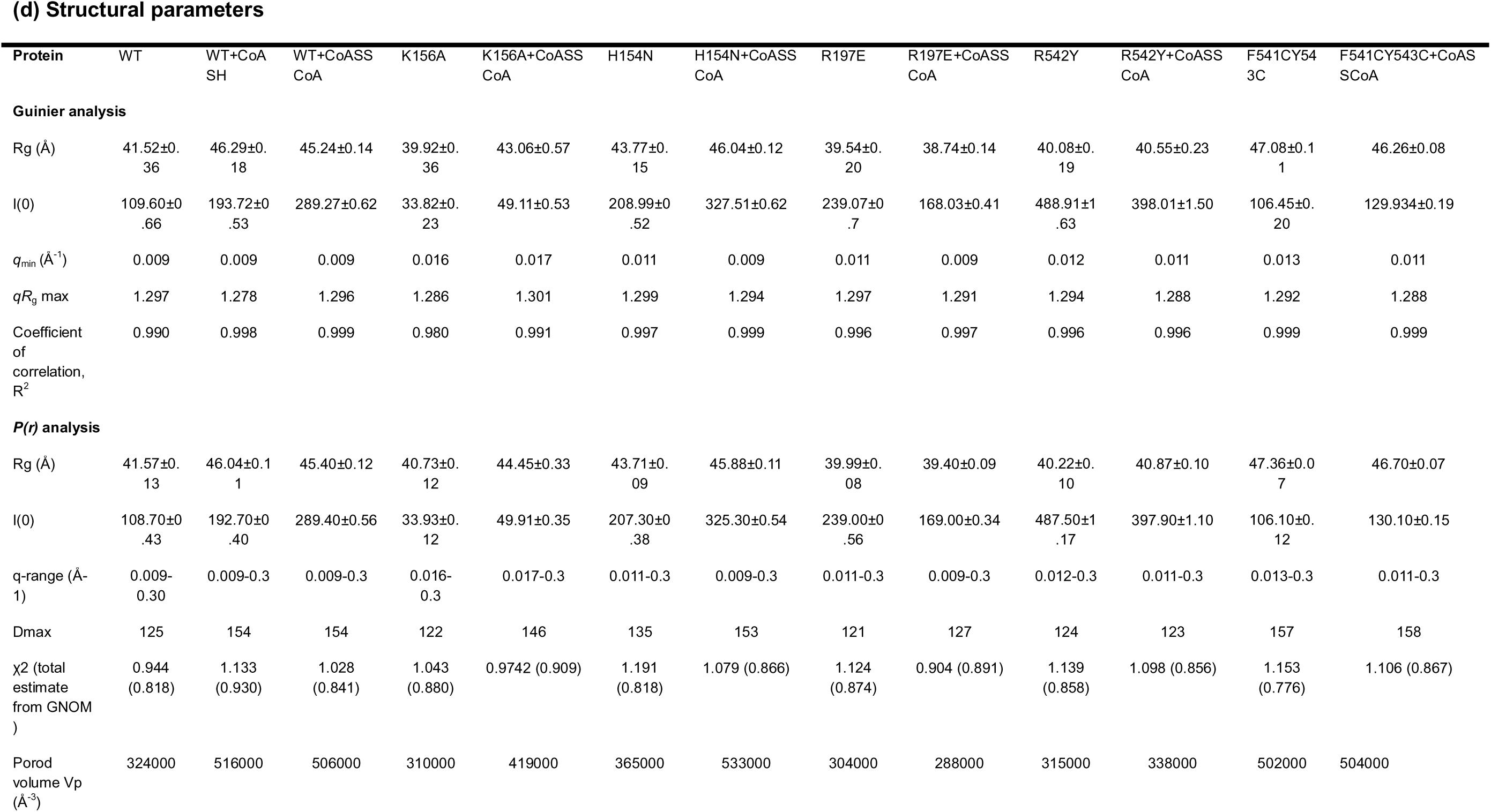
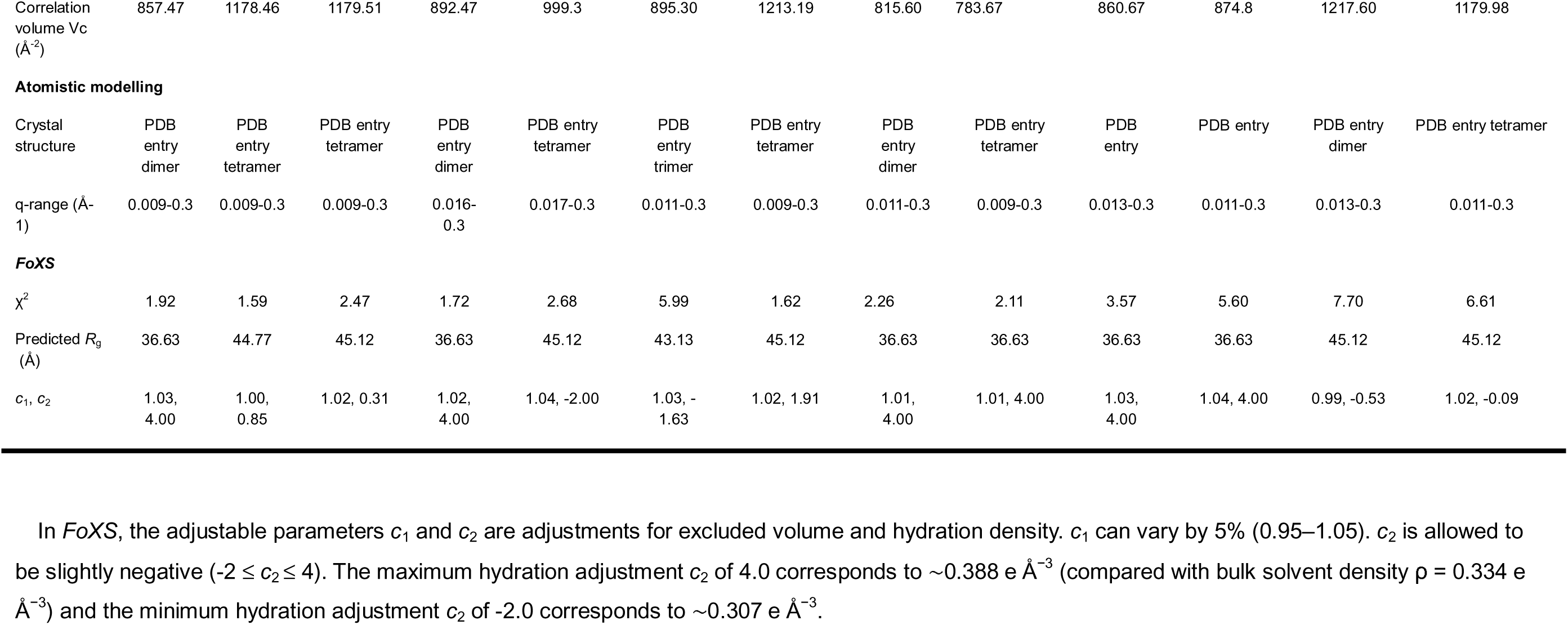
Statistics of SAXS analysis.

**(a) Sample details.**
|  |  |
| --- | --- |
| Protein | ME2 |
| Organism | *Homo sapiens (Human)* |
| Source | *E. coli* expressed |
| UniProt sequence ID | P23368 |
| (residues in construct) | (18-584) |
| Extinction coefficient [$A_{280}$, 0.1%(w/v)] | 0.883 |
| Molecular mass $M$ from chemical composition (kDa) | 63.6 |
| concentration (mg ml<sup>-1</sup>) | 1.0 |
| $\bar{v}$ from chemical composition (cm<sup>3</sup> g<sup>-1</sup>) | 0.742 |
| Particle contrast from sequence and solvent constituents, $\Delta\bar{\rho}$ ($\rho_{\text{protein}} - \rho_{\text{solvent}}$; 10<sup>10</sup> cm<sup>-2</sup>) | 2.76 (12.23-9.47) |
| Solvent | 20 mM Tris-HCl pH 7.4, 150 mM KCl, 1 mM DTT |
| **(b) SAXS data-collection parameters** |  |
| Instrument/data processing | BL19U2 beamline at the Shanghai Synchrotron Radiation Facility (SSRF) with Dectris PILATUS 1M detector |
| Wavelength (Å) | 1.03 |
| Beam size (μm) | 380 (H) ×25 (V) |
| Camera length (m) | 2.683 |
| $q$-measurement range (Å<sup>-1</sup>) | 0.008~0.45 |
| Absolute scaling method | Comparison with scattering from 1 mm pure H<sub>2</sub>O |
| Normalization | To transmitted intensity by beam-stop counter |
| Monitoring for radiation damage | X-ray dose maintained below 210 Gy, data frame-by-frame comparison |
| Exposure time | Continuous 1 s data-frame measurements |
| Sample temperature (°C) | 10.0 |
| **(c) Software employed for SAXS data reduction, analysis, and interpretation.** |  |
| SAXS data reduction | *RAW (Hopkins et al., 2017)* |
| Extinction coefficient estimate | *ProtParam (Gasteiger et al., 2005)* |
| Calculation of $\Delta\bar{\rho}$ and $\bar{v}$ values | *MULCh 1.1 (Whitten et al., 2008)* |
| Basic analyses: Guinier, $P(r)$, $V_p$ | *RAW (Hopkins et al., 2017)* |
| Atomic structure modelling | *FoXS (Schneidman-Duhovny et al., 2013)* |

(d) Structural parameters
| Protein | WT | WT+CoA<br>SH | WT+CoASS<br>CoA | K156A | K156A+CoASS<br>CoA | H154N | H154N+CoASS<br>CoA | R197E | R197E+CoASS<br>CoA | R542Y | R542Y+CoASS<br>CoA | F541CY54<br>3C | F541CY543C+CoAS<br>SCoA |
| --- | --- | --- | --- | --- | --- | --- | --- | --- | --- | --- | --- | --- | --- |
| **Guinier analysis** |  |  |  |  |  |  |  |  |  |  |  |  |  |
| R<sub>g</sub> (Å) | 41.52±0.36 | 46.29±0.18 | 45.24±0.14 | 39.92±0.36 | 43.06±0.57 | 43.77±0.15 | 46.04±0.12 | 39.54±0.20 | 38.74±0.14 | 40.08±0.19 | 40.55±0.23 | 47.08±0.11 | 46.26±0.08 |
| I(0) | 109.60±0.66 | 193.72±0.53 | 289.27±0.62 | 33.82±0.23 | 49.11±0.53 | 208.99±0.52 | 327.51±0.62 | 239.07±0.7 | 168.03±0.41 | 488.91±1.63 | 398.01±1.50 | 106.45±0.20 | 129.934±0.19 |
| q<sub>min</sub> (Å<sup>-1</sup>) | 0.009 | 0.009 | 0.009 | 0.016 | 0.017 | 0.011 | 0.009 | 0.011 | 0.009 | 0.012 | 0.011 | 0.013 | 0.011 |
| qR<sub>g</sub> max | 1.297 | 1.278 | 1.296 | 1.286 | 1.301 | 1.299 | 1.294 | 1.297 | 1.291 | 1.294 | 1.288 | 1.292 | 1.288 |
| Coefficient of correlation, R<sup>2</sup> | 0.990 | 0.998 | 0.999 | 0.980 | 0.991 | 0.997 | 0.999 | 0.996 | 0.997 | 0.996 | 0.996 | 0.999 | 0.999 |
| **P(r) analysis** |  |  |  |  |  |  |  |  |  |  |  |  |  |
| R<sub>g</sub> (Å) | 41.57±0.13 | 46.04±0.11 | 45.40±0.12 | 40.73±0.12 | 44.45±0.33 | 43.71±0.09 | 45.88±0.11 | 39.99±0.08 | 39.40±0.09 | 40.22±0.10 | 40.87±0.10 | 47.36±0.07 | 46.70±0.07 |
| I(0) | 108.70±0.43 | 192.70±0.40 | 289.40±0.56 | 33.93±0.12 | 49.91±0.35 | 207.30±0.38 | 325.30±0.54 | 239.00±0.56 | 169.00±0.34 | 487.50±1.17 | 397.90±1.10 | 106.10±0.12 | 130.10±0.15 |
| q-range (Å<sup>-1</sup>) | 0.009-0.30 | 0.009-0.3 | 0.009-0.3 | 0.016-0.3 | 0.017-0.3 | 0.011-0.3 | 0.009-0.3 | 0.011-0.3 | 0.009-0.3 | 0.012-0.3 | 0.011-0.3 | 0.013-0.3 | 0.011-0.3 |
| Dmax | 125 | 154 | 154 | 122 | 146 | 135 | 153 | 121 | 127 | 124 | 123 | 157 | 158 |
| χ<sup>2</sup> (total estimate from GNOM) | 0.944 (0.818) | 1.133 (0.930) | 1.028 (0.841) | 1.043 (0.880) | 0.9742 (0.909) | 1.191 (0.818) | 1.079 (0.866) | 1.124 (0.874) | 0.904 (0.891) | 1.139 (0.858) | 1.098 (0.856) | 1.153 (0.776) | 1.106 (0.867) |
| Porod volume V<sub>p</sub> (Å<sup>3</sup>) | 324000 | 516000 | 506000 | 310000 | 419000 | 365000 | 533000 | 304000 | 288000 | 315000 | 338000 | 502000 | 504000 |
| Correlation volume $V_c$ ($\text{\AA}^{-2}$) | 857.47 | 1178.46 | 1179.51 | 892.47 | 999.3 | 895.30 | 1213.19 | 815.60 | 783.67 | 860.67 | 874.8 | 1217.60 | 1179.98 |
| **Atomistic modelling** |  |  |  |  |  |  |  |  |  |  |  |  |  |
| Crystal structure | PDB entry dimer | PDB entry tetramer | PDB entry tetramer | PDB entry dimer | PDB entry tetramer | PDB entry trimer | PDB entry tetramer | PDB entry dimer | PDB entry tetramer | PDB entry | PDB entry | PDB entry dimer | PDB entry tetramer |
| q-range ($\text{\AA}^{-1}$) | 0.009-0.3 | 0.009-0.3 | 0.009-0.3 | 0.016-0.3 | 0.017-0.3 | 0.011-0.3 | 0.009-0.3 | 0.011-0.3 | 0.009-0.3 | 0.013-0.3 | 0.011-0.3 | 0.013-0.3 | 0.011-0.3 |
| **FoXS** |  |  |  |  |  |  |  |  |  |  |  |  |  |
| $\chi^2$ | 1.92 | 1.59 | 2.47 | 1.72 | 2.68 | 5.99 | 1.62 | 2.26 | 2.11 | 3.57 | 5.60 | 7.70 | 6.61 |
| Predicted $R_g$ ($\text{\AA}$) | 36.63 | 44.77 | 45.12 | 36.63 | 45.12 | 43.13 | 45.12 | 36.63 | 36.63 | 36.63 | 36.63 | 45.12 | 45.12 |
| $c_1, c_2$ | 1.03, 4.00 | 1.00, 0.85 | 1.02, 0.31 | 1.02, 4.00 | 1.04, -2.00 | 1.03, -1.63 | 1.02, 1.91 | 1.01, 4.00 | 1.01, 4.00 | 1.03, 4.00 | 1.04, 4.00 | 0.99, -0.53 | 1.02, -0.09 |

In *FoXS*, the adjustable parameters $c_1$ and $c_2$ are adjustments for excluded volume and hydration density. $c_1$ can vary by 5% (0.95–1.05). $c_2$ is allowed to be slightly negative ( $-2 \leq c_2 \leq 4$ ). The maximum hydration adjustment $c_2$ of 4.0 corresponds to $\sim 0.388 \text{ e \AA}^{-3}$ (compared with bulk solvent density $\rho = 0.334 \text{ e \AA}^{-3}$ ) and the minimum hydration adjustment $c_2$ of -2.0 corresponds to $\sim 0.307 \text{ e \AA}^{-3}$ .

**Supplementary Table 2.** Data collection and refinement statistics of ME2.

|  | Apo<br>(PDB 25EH) | CoASH<br>(PDB 25EI) | CoA disulfide<br>(PDB 25EJ) | CoASH/NAD<sup>+</sup>/P<br>YR/Mn<sup>2+</sup><br>(PDB 25EK) | CoA<br>disulfide/NAD<sup>+</sup>/<br>PYR/Mn<sup>2+</sup><br>(PDB 25EL) |
| --- | --- | --- | --- | --- | --- |
| Data collection |  |  |  |  |  |
| Wavelength | 0.9792 | 0.9778 | 0.9792 | 0.9792 | 0.9792 |
| Space group | *P2*<sub>1</sub> | *P2*<sub>1</sub> | *C2* | *C2* | *C2* |
| Cell<br>dimensions |  |  |  |  |  |
| a,b,c (Å) | 107.94, 113.29,<br>109.43 | 106.77, 116.89,<br>106.94 | 197.26, 79.10,<br>169.029 | 204.44, 59.13,<br>106.91 | 203.88, 57.81,<br>144.49 |
| α,β,γ (°) | 90.00, 103.31,<br>90.00 | 90.00, 98.30,<br>90.00 | 90.00, 117.59,<br>90.00 | 90.00, 101.83,<br>90.00 | 90.00, 134.29,<br>90.00 |
| Resolution<br>(Å)<sup>†</sup> | 50.00-2.50<br>(2.54-2.50) | 50.00-2.50<br>(2.54-2.50) | 50.00-2.90<br>(2.95-2.90) | 52.32-2.13<br>(2.25-2.13) | 103.43-2.32<br>(2.38-2.32) |
| R<sub>sym</sub> (%) | 11.5 | 9.8 | 5.4 | 7.7 | 6.8 |
| I/σ | 15.20 (3.30) | 11.58 (2.45) | 20.47 (2.08) | 13.6 (2.5) | 11.2 (2.1) |
| Completeness<br>(%) | 90.3 (57.5) | 92.0 (79.6) | 98.7 (99.0) | 95.4 (87.1) | 100.0 (99.9) |
| Total No. of<br>reflections | 3871391 | 3208240 | 721285 | 451795 | 356132 |
| Unique<br>reflections | 81228 | 81419 | 50544 | 66939 | 52652 |
| Redundancy | 5.3 (2.9) | 5.6 (4.5) | 3.3 (3.3) | 6.7 (6.7) | 6.8 (6.3) |
| Refinement |  |  |  |  |  |
| Resolution (Å) | 48.24-2.48<br>(2.54-2.48) | 48.19-2.51<br>(2.57-2.51) | 42.26-2.90<br>(2.98-2.90) | 50.01-2.13<br>(2.19-2.13) | 50.01-2.32<br>(2.38-2.32) |
| No. of<br>reflections | 75409 (3929) | 72351 (3981) | 44814 (2311) | 63571 (3364) | 50232 (2393) |
| R<sub>work</sub>/R<sub>free</sub>(%) | 22.13/24.19 | 21.57/23.44 | 19.08/22.72 | 21.00/23.61 | 21.64/24.50 |
| No. Of atoms |  |  |  |  |  |
| Protein | 17070 | 17418 | 16932 | 8576 | 8657 |
| Ligand | 5 | 232 | 192 | 199 | 220 |
| Water | 435 | 257 | 35 | 69 | 112 |
| *B*-factors(Å<sup>2</sup>) |  |  |  |  |  |
| Protein | 35.84 | 41.58 | 56.56 | 54.45 | 66.70 |
| Ligand | 31.31 | 36.76 | 50.95 | 55.73 | 67.59 |
| Water | 30.67 | 30.29 | 38.48 | 46.25 | 60.79 |
| rms deviations |  |  |  |  |  |
| Bond lengths (Å) | 0.003 | 0.003 | 0.004 | 0.004 | 0.004 |
| Bond angles (°) | 1.262 | 1.272 | 1.362 | 1.309 | 1.300 |
| Ramachandra n Plot (%)<sup>1</sup> | 97.1/2.8/0.1 | 96.5/3.4/0 | 95.6/4.3/0.1 | 97.1/2.9/0 | 96.8/3.1/0.1 |

Three crystal experiments for each structure.
$R_{sym} = \frac{\sum_h \sum_i |I_{h,i} - I_h|}{\sum_h \sum_i I_{h,i}}$ , where $I_h$ is the mean intensity of the $i$ observations of symmetry related reflections
of $h$ .
<sup>1</sup>Residues in favored, allowed, and outlier regions of the Ramachandran plot.

